# KIRA6 is an effective and versatile mast cell inhibitor of IgE-mediated activation

**DOI:** 10.1101/2024.04.18.589885

**Authors:** Veronika Wunderle, Thomas Wilhelm, Shatha Boukeileh, Jonas Goßen, Michael A. Margreiter, Roman Sakurov, Sandro Capellmann, Maike Schwoerer, Nabil Ahmed, Gina Bronneberg, Michel Arock, Christian Martin, Thomas Schubert, Francesca Levi-Schaffer, Giulia Rosetti, Boaz Tirosh, Michael Huber

## Abstract

Incidents of IgE-mediated, mast cell (MC)-driven allergic diseases are constantly rising and there is an urgent need for the development of novel pharmacological MC stabilizers. Allergen/antigen (Ag)-triggered activation of MCs via crosslinking of the high-affinity receptor for IgE (FcεRI) is regulated, amongst others, by the coordinated action of various cytosolic tyrosine kinases of the SRC family, e.g. LYN and FYN, which exert positive as well as negative functions. We report that KIRA6, an inhibitor developed for the endoplasmic reticulum (ER) stress sensor IRE1α, suppresses IgE-mediated pro-inflammatory MC activation by inhibiting both LYN and FYN. KIRA6 dose-dependently and effectively attenuates Ag-stimulated early signaling (e.g. substrate tyrosine phosphorylation, Ca^2+^ mobilization, and activation of MAPK pathways) as well as effector functions such as degranulation and pro-inflammatory cytokine production/secretion in murine bone marrow-derived MCs (BMMCs). Moreover, Ag-triggered bronchoconstriction in an *ex vivo* model of precision-cut lung slices (PCLS), and IgE-mediated stimulation of human MCs were repressed by KIRA6. To get in-depth inside into KIRA6 interaction with three MC-relevant tyrosine kinases, LYN, FYN, and KIT, and to elicit the potential of KIRA6 structure to serve as pharmacophore for the development of respective single-, dual-, or triple-specificity inhibitors, we modeled and evaluated the binding of KIRA6 on the three kinases by applying homology modeling and molecular dynamics simulations, as well as MM GBSA calculations. We found that KIRA6 has a high propensity to bind the inactive state of LYN, FYN, and KIT with comparable affinities. In conclusion, our data suggest the use of novel inhibitors based on the KIRA6 pharmacophore as effective MC stabilizers to improve treatment of pro-inflammatory diseases with MC involvement in need of effective pharmacological interventions.

## Introduction

Pharmacological modulation of enzymatic activities of signaling proteins, such as protein and lipid kinases, has become general practice in the treatment of inflammatory and neoplastic diseases (Akinleye et al., 2013)(Marone et al., 2008). Particularly in the latter, cells frequently develop resistance to the drug used, amongst others, by directly acquiring mutations in the drug target rendering it inaccessible to the drug, or by the induction/activation of additional signaling proteins leading to the re-structuring of whole signaling pathways (Johannessen et al., 2010) (Falchi & Verstovsek, 2018). Consequently, a second or third drug should be administered to regain treatment success. Drugs with the potential to inhibit different enzymes, preferentially in structurally or kinetically different pathways, should reduce the development of resistance mechanisms. Similar points would account for inflammatory diseases, e.g. allergic hypersensitivity diseases. Here, not the development of drug-resistant cells, but the insufficiency of only one drug to control different aspects of the activation of one or more cell types involved would pose limitations. Especially in allergies or other mast cell (MC)-mediated diseases, the combined blockade of initial, receptor-proximal activation processes as well as the kinetically downstream production of a multitude of mediators, such as cytokines and chemokines, with the potential to activate further inflammatory cells, would be desirable.

A central mechanism of allergic MC activation is initiated by the antigen/allergen (Ag)-mediated cross-linking of the IgE-loaded high-affinity receptor for IgE (FcεRI) and subsequent induction of various signaling pathways eventually resulting in typical MC responses such as degranulation (release of pre-formed mediators from secretory lysosomes) and production of lipid mediators as well as pro-inflammatory cytokines (e.g. IL-6 and TNF) (Gilfillan & Rivera, 2009)). A crucial signaling enzyme in FcεRI-mediated MC activation is the SRC family kinase (SFK) LYN, which constitutively interacts with the FcεRI β-chain (Vonakis et al., 1997). After receptor cross-linking, LYN phosphorylates the tyrosine (Tyr) residues in the immunoreceptor Tyr-based activation motives (ITAMs) of the FcεRI β-and γ-chains, allowing for consequent interaction with SH2-domain-containing proteins thus initiating differential signaling pathways necessary for MC activation, e.g. phospholipase Cγ (PLCγ), phosphatidylinositol-3-kinase (PI3K), and mitogen-activated protein kinases (MAPKs) controlled processes (Gilfillan & Rivera, 2009).

Once PLCγ is recruited to the plasma membrane, particularly by the transmembrane adaptor protein LAT1 (Linker for activated T cells 1), it hydrolyzes its substrate phosphatidylinositol- 4,5-bisphosphate (PI(4,5)P_2_) to yield inositol-1,4,5-trisphosphate (IP_3_) and diacylglycerol (DAG) (Gilfillan & Rivera, 2009)). Whereas DAG promotes activation of conventional and novel isotypes of PKC, IP_3_ binds to its receptor in the membrane of the endoplasmic reticulum (ER), which results in the release of Ca^2^ from the ER and eventually in the STIM1/ORAI1-controlled influx of extracellular Ca(Baba et al., 2008). Both Ca^2+^ influx and PKC activation are mandatory for the process of degranulation (Sagi-Eisenberg et al., 1985)(Cochrane & Douglas, 1974). PI3K and MAPKs are particularly involved in transcriptional processes relevant for the production of pro-inflammatory cytokines (Gilfillan & Rivera, 2009). Inhibition of signaling processes in the course of allergic MC activation, preferably close to the initial activating event (e.g. LYN activation), would be desirable. However, assigning only activating or inhibitory functions to a protein is hard in the context of cellular signaling, since activators can be initiators of negative feedback mechanisms as well (Huber, 2013). LYN, for instance, can phosphorylate and activate the lipid phosphatase SH2-domain-containing inositol 5’-phosphatase 1 (SHIP1), a dominant negative regulator of the PI3K pathway, which hydrolyzes the PI3K-generated phospholipid phosphatidylinositol-3,4,5-trisphosphate (PIP(HernandezLJHansen et al., 2004)(Miranda et al., 2016). Moreover, LYN is engaged in the attenuation of further SFKs, such as FYN, by contributing to the activation of the C-terminal SRC kinase (CSK), which then phosphorylates SFKs at their inhibitory, C-terminal Tyr residue allowing for intramolecular SH2-domain/phospho-Tyr interaction (Gilfillan & Rivera, 2009).

Both proliferating and secreting cells are particularly dependent on ER-mediated translation of proteins involving accurate folding and assembly of effective complexes while still in the lumen of the ER. An elaborate control system has evolved, called the unfolded protein response (UPR), which detects unfolded/misfolded proteins and activates cellular mechanisms to relieve folding stress (adaptive UPR) (Grootjans et al., 2016). Three sensor proteins in the membrane of the ER detect such unfolded proteins hence resulting in the activation of sensor-specific transcription factors that allow adaptation to this cellular stress by activating genes involved in ER expansion, protein folding, and aggregate degradation, amongst others (Hetz et al., 2020). These sensors are inositol-requiring 1 enzyme α (IRE1α), protein kinase RNA-like ER kinase (PERK), and activating transcription factor 6 (ATF6), with IRE1α being the evolutionary oldest. IRE1α comprises a luminal N-terminal binding immunoglobulin protein/heat shock protein 5 (BIP/HSPA5)-binding domain, a central cytoplasmic serine/threonine (Ser/Thr) kinase domain and a C-terminal ribonuclease (RNase) domain. The presence of misfolded proteins causes the chaperone Binding Immunoglobulin Protein (BIP) to dissociate from IRE1α causing its oligomerization, autophosphorylation, and activation of the RNase domain (Hetz et al., 2020). Active IRE1α RNase then non-conventionally splices the mRNA of the transcription factor X-box binding protein 1 (XBP1), allowing its stable translation, nuclear translocation, and transcription of genes involved in adaptive UPR. Excess folding stress overburdens the adaptive UPR and switches it, promoted by the stress sensor PERK, to a terminal response (apoptosis) (Hetz et al., 2020).

The development of various small molecule inhibitors against the three UPR sensors has allowed unravelling differential functions of the UPR (Maly & Papa, 2014). Moreover, the dependence on an effective UPR system of tumor cells holds great promise for the use of such inhibitors in cancer therapy. In recent years, however, cross-reactive proteins of certain UPR inhibitors were identified, which were even more sensitive to the respective inhibitor than the initial target (Mahameed et al., 2019)(Rojas-Rivera et al., 2017). In this study, we have found that the IRE1α inhibitor KIRA6 (kinase inhibitor and RNase attenuator 6)(Ghosh et al., 2014) is a potent inhibitor of the SFKs LYN and FYN, thus effectively blocking IgE-mediated MC activation. In-depth molecular modeling analysis revealed enhanced affinity of KIRA6 for the inactive conformations of the catalytic centers of LYN, FYN, and KIT, and thus offers a promising pharmacophore on the basis of which forceful MC-stabilizing, anti-allergy remedies can be developed.

## Material and Methods

### Cell culture, cell lines and IgE sensitization

Murine bone marrow-derived mast cells (BMMCs) from WT or mutant mice deficient for SHIP1 (gene name: Inpp5d) (*Ship1^-/-^*) or LYN (*Lyn^-/-^*) were generated as described previously (Meurer et al., 2016) from 6- to 8-week-old mice (littermates, mixed C57BL/6x129/Sv background).

Hematopoietic bone marrow stem cells were cultured in RPMI 1640 medium + L-glutamine (Thermo Fisher Scientific, #21875-034) containing 1% X63Ag8-653-conditioned medium (source of IL-3), 15% FCS (Capricorn, #FCS-12A), 10 mM HEPES (Sigma-Aldrich, #H0887), 100 µM β-mercaptoethanol (Sigma-Aldrich, #M6250), 100 units/ml penicillin and 100 µg/ml streptomycin (Sigma-Aldrich, #P0781). After a differentiation period of 4 – 6 weeks, > 95% of the cells were KIT- and FcεRI-positive confirming successful MC differentiation.

BMMCs were sensitized for Ag stimulation with dinitrophenol (DNP) coupled to human serum albumin (HSA) (DNP-HSA, Sigma-Aldrich, #A6661) by pre-loading the FcεRI with 0.15 µg/ml IgE (clone SPE-7, Sigma-Aldrich, #A2831) overnight. If starvation was indicated, the culture medium was replaced by medium containing 10% FCS and lacking IL-3, 12 - 16 h before the experiment.

All experiments were performed in accordance with German legislation governing animal studies and following the Principles of Laboratory Animal Care. Mice are held in the Institute of Laboratory Animal Science, Medical Faculty of RWTH Aachen University, which holds a license for husbandry and breeding of laboratory animals from the Veterinary Office of the Staedteregion Aachen (Administrative District). The Institute follows a Quality Management System, which is certified according to DIN ISO 9001/2015. Every step in this project involving mice was reviewed by the animal welfare officer. All experiments were approved by the Landesamt fulr Natur, Umwelt und Verbraucherschutz NRW (LANUV), Recklinghausen (AZ 84-02.04.2016.A496).

The mast cell leukemia (MCL) cell line HMC 1.2 (KIT^V560G, D816V^), kindly provided by Dr. J. Butterfield (Mayo Clinic) (Sundström et al., 2003) was cultured in RPMI-1640 medium (Thermo Fisher Scientific, #21875-034) supplemented with 10% FCS (Capricorn, #FCS-12A) and 100 units/ml penicillin + 100 µg/ml streptomycin (Sigma-Aldrich, #P0781) in a humified atmosphere containing 5% CO_2_. The medium was renewed twice a week.

ROSA^KIT^ ^WT^ cells were cultured as previously described (Saleh et al., 2014) in IMDM medium (Thermo Fisher Scientific, #21980-032) supplemented with 100 units penicillin/ml and 100 µg/ml streptomycin (Sigma-Aldrich, #P0781), 1 mM sodium pyruvate (Thermo Fisher Scientific , #11360-039), MEM vitamin solution (Thermo Fisher Scientific , #11120-037), MEM amino acids (Thermo Fisher Scientific, #11130-036), 2 mM L-glutamine (Thermo Fisher Scientific, #25030-024), Insulin-Transferrin-Selenium (Thermo Fisher Scientific, #41400-045), 100 µM β-mercaptoethanol (Sigma-Aldrich, #M6250), 0.1% BSA (Serva, #11930.04) and 80 ng/ml mSCF from CHO culture supernatant (Malbec et al., 2007). ROSA^KIT^ ^WT^ cells were primed for 4 days with rh-IL-4 (20 ng/ml; ImmunoTools, #1134B0045) and human monoclonal IgE (2 µg/ml; Sigma-Aldrich, #401152) to enhance the expression of the FcεRI before stimulation (Saleh et al., 2014).

Human cord blood-derived MCs (CBMCs) were obtained by culturing umbilical cord blood mononuclear cells as previously described (Suzuki et al., 1997). In brief, fresh cord blood diluted in Hank’s solution was loaded on Lymphoprep (Serumwerk Bernburg, #1858) and centrifuged (350 x g, 25 min, 24 °C). Mononuclear cells were washed twice and resuspended in a minimum essential medium containing 10 µg/mL ribonucleases (MEM-α nucleosides; Gibco, #12571063) supplemented with 10% heat-inactivated FCS (HyClone, GE Healthcare, SV30160.03), 100 U/mL penicillin and 100 µg/mL streptomycin (Gibco, #15140-122) and containing recombinant human stem cell factor (h-SCF; Gibco, PeproTech, #300-07; 100 ng/mL) and human IL-6 (Gibco, PeproTech, #200-06; 20 ng/mL). Prostaglandin E2 (PGE2; Sigma-Aldrich, #P0409; 1 ng/mL) was also regularly added in order to prevent any monocytic development. Cells were maintained in a humidified incubator (37 °C, 5% CO_2_) with media replaced on a weekly basis. Following a 7 - 8 week culture period, CBMCs were used following examination for maturity and viability. Cord blood was obtained according to the Institutional Helsinki Committee guidelines of Hadassah Hospital, and its use was approved by the committee.

For activation of CBMCs, human myeloma IgE (0.3 µg/mL, Sigma-Aldrich, #401152) was used for sensitizing the cells in the presence of recombinant human IL-4 (Gibco, PeproTech, #200-04; 10 ng/mL) for three days. Subsequently, cells were washed twice and resuspended in Tyrode’s buffer (consisting of 137 mM NaCl, 5.5 mM glucose, 2 mM KCl, 12 mM NaHCO_3_, and 0.3 mM Na_2_HPO_4_ and supplemented with 1.8 mM CaCl_2_ and 0.9 mM MgCl_2_; pH 7.34 for a short stimulation of 30 min) or in a CBMC medium supplemented with recombinant h-SCF (100 ng/mL, Gibco, PeproTech, #300-07) (for a longer stimulation). CBMCs were activated with polyclonal rabbit anti-human IgE Ab (5 µg/mL, Dako, #A0094).

## Materials

KIRA6 was purchased from Cayman Chemical (#19151). KIRA8 (#SML2903), A23187 (#C7522), Dinitrophenol-human serum albumin (DNP-HSA, #A6661) and monoclonal IgE with specificity for DNP (clone Spe-7, Sigma, #A2831) were from Sigma Aldrich. Murine stem cell factor (SCF) was from PeproTech (#250-03). PMA (#1585) came from Sigma-Aldrich and Thapsigargin (TG, #ab147487) from Abcam. Saracatinib (#S1006) was purchased from Selleckchem and Tunicamycin (#A2242) from Applichem. DMSO (#4720.1) was obtained from Carl Roth GmbH & Co.

### Calcium measurement

BMMCs were pre-loaded with IgE and starved (10% FCS, no IL-3) overnight. Cells were resuspended at a density of 1 x 10^7^ cells/ml in RPMI 1640 containing 1% FCS, 0.1% BSA (Serva, #11930.04), 1.3 μM Fluo-3 AM (#F1241), 2.7 μM Fura Red AM (#F3020) and 0.1% pluoronic F-127 (all ThermoFisher) and incubated for 45 min at 37°C for flow cytometry analysis in a FACSCalibur flow cytometer (BD Biosciences). Steady-state fluorescence was measured for 1 min before 20 ng/ml DNP-HSA were added for 4 min. Subsequently, the ratios of the Ca^2+^-bound /Ca^2+^-unbound were used to generate line graphs from FACS profiles using the FlowJo analysis software (Treestar) and normalized areas under the curve (AUC) were calculated for objectification.

### ELISAs

To determine IL-6 and TNF secretion, murine BMMCs were stimulated as indicated in the respective experiments. If cells were stimulated with Ag, MCs were pre-loaded with IgE (clone Spe-7, 0.15 µg/ml). Cell number was adjusted to 1.2 x 10^6^ cells/ml in stimulation medium (RPMI 1640 + 0.1% BSA (Serva, #11930.04), cells were allowed to adapt to 37°C and stimulated for 3 h. Supernatants were collected to determine cytokine release. 96-well ELISA plates (Corning, #9018) were coated with capturing anti-IL-6 (1:250, BD Biosciences, #554400) or anti-TNF (1:200, R&D Systems, #AF410-NA) antibodies diluted in PBS overnight at 4°C according to manufacturer’s instructions. ELISA plates were washed three times with PBS+0.1% Tween and blocked with PBS+2% BSA (IL-6 ELISA) or PBS+1%BSA+5% sucrose (TNF ELISA) before loading of supernatants (50 µl for IL-6 ELISA, 100 µl for TNF ELISA). Additionally to loading of supernatants, IL-6 (BD Pharmingen, #554582) and TNF (R&D Systems, #410-MT-010) serial 1:2 standard dilutions were added and plates were incubated overnight at 4°C. Thereupon, plates were washed three times again followed by incubation with biotinylated anti-IL-6 (1:500, BD Biosciences, #554402) and anti-TNF (1:250, R&D Systems, #BAF-410) antibodies diluted in PBS+1% BSA for 45 min and 2 h, respectively, at room temperature (RT). After 3 washing steps, streptavidin alkaline phosphatase (SAP, 1:1000, BD Pharmingen, #554065) was added for 30 min at RT. After 3 more washing steps, the substrate *p*-Nitro-phenyl-phosphate (1 pill per 5 ml in sodium carbonate buffer (2mM MgCl_2_ in 50 mM sodium carbonate, pH 9.8), (Sigma, #S0942-200TAB) was added and OD_450_ was recorded using a plate reader (BioTek Eon). Qualitative differences and similarities between WT and mutant cells were consistent throughout the study. Levels of secreted cytokines varied due to batch-to-batch variations of primary, differentiated cells of different age and from different mice.

### RT-qPCR

RNA was extracted from 4 x 10^6^ cells using RNeasy Mini Kit (Qiagen, #74106) according to the manufacturer’s instructions. 1 µg RNA was used for reverse transcription using random hexamer primer (Roche, #11034731001) and Omniscript Kit (Qiagen, #205113) as instructed by the manufacturer. Quantitative PCR was performed on a Rotorgene (Qiagen) using the Sybr green reaction mix (Meridian Bioscience, #QT650-05). Expression was normalized to the reference gene *Hprt/HPRT*. Relative expression ratios were calculated according to the ddC_t_ method (Pfaffl, 2001).

Primer sequences were as follows: *Hprt* fwd GCT GGT GAA AAG GAC CTC T, *Hprt* rev CAC AGG ACT AGA ACA CCT GC; *Il6* fwd TCC AGT TGC CTT CTT GGG AC, *Il6* rev GTG TAA TTA AGC CTC CGA CTT G; *Xbp1s* fwd AAG AAC ACG CTT GGG AAT GG, *Xbp1s* rev CTG CAC CTG CTG CGG Ac.

*human IL8* fwd CAC TGC GCC AAC ACA GAA AT, *IL8* rev ATG AAT TCT CAG CCC TCT TCA A; human *HPRT* fwd TGA CAC TGG CAA AAC AAT GCA, *HPRT* rev GGT CCT TTT CAC CAG CAA GCT; human *XBP1s fwd AAC CAG GAG TTA AGA CAG CGC TT, XBP1s rev CTG CAC CTG CTG CGG ACT*

### Degranulation assays (**β**-hexosaminidase and LAMP1 assay)

To measure degranulation using release of β-hexosaminidase as a readout, non-starved IgE pre-loaded BMMCs were washed in sterile PBS, resuspended in Tyrode’s buffer (130 mM NaCl, 5 mM KCl, 1.4 mM CaCl_2_, 1 mM MgCl_2_, 5.6 mM glucose and 0.1% BSA in 10 mM Hepes, pH 7.4) at a density of 1.2 x 10^6^, adapted to 37°C and stimulated as indicated. The supernatant was collected by centrifugation and the remaining cell pellet was lysed in Tyrode’s buffer containing 10% NP-40. 10 µl of both, pellet and supernatant, were incubated with 50 µl of substrate solution (1.3 mg/ml *p*-Nitrophenyl-N-acetyl-β-D-glucose in 0.1 M sodium citrate, pH 4.5).

The measurement of *p*-nitrophenol, generated by β-hexosaminidase, was done by a spectrophotometric reader (BioTek^®^ Eon^TM^) at a wavelength of λ = 405 nm (Huber et al., 1998). The amount of degranulation in percent was determined as follows:

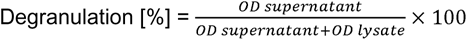

For LAMP1 assay, IgE pre-loaded BMMCs were washed in sterile PBS, resuspended in RPMI 1640 containing 0.1% BSA and adapted to 37°C. Next, the indicated treatment was performed, cells were pelleted, washed in FACS buffer (PBS, 3% FCS, 0.1% sodium azide) and stained with FITC-conjugated anti-LAMP1 antibody (CD107a) (BioLegend, #121605) for 25 min at 4°C. LAMP1 externalization was determined by flow cytometry. Data were analyzed and mean fluorescence intensities (MFIs) were calculated using the FlowJo analysis software.

### Immunoprecipitation and Western blotting

20 x 10^6^ BMMCs per condition were starved and pre-loaded with IgE overnight, washed with PBS, resuspended in RPMI + 0.1% BSA at a density of 10 x 10^6^ and allowed to adapt to 37°C. Treatment was performed as indicated in the respective figures and figure legends. Cells were then pelleted and lysed in phosphorylation solubilization buffer (PSB; 0.5% IGEPAL, 1 mM Na_3_VO_4_, 0.5% sodium desoxycholate and protease inhibitors (10 µg/ml aprotinin, 4 mg/ml leupeptin, 1mM PMSF) for 1 h at 4°C. After normalization for protein content (Pierce BCA Protein Assay Kits, #23227), samples were mixed with Laemmli buffer and boiled for 5 min at 95°C, whole-cell lysates were subjected to SDS-PAGE and subsequent Western blot (WB) analysis (Gimborn et al., 2005).

After protein transfer, the polyvinylidenfluorid (PVDF) membrane was dried, hydrophilized by methanol and blocked in PBST (PBS + 0.5% Tween) wash buffer with dry non-fat milk powder (5% milk powder (w/v) in PBS + 0.5% Tween) for 30 min, before incubation with primary antibody diluted according to manufacturer’s instructions overnight at 4°C. Thereupon, membranes were washed three times and incubated with horseradish peroxidase (HRP)-coupled secondary antibody diluted in PBS + 0.5% Tween, the membrane was developed with enhanced chemiluminescence solution using a LAS 4000mini (FujiFilm). The expression of p85, HSP90 or GAPDH served as control for comparable loading. Since the stimulation times were short (less than 15 min), differences in expression between these loading controls and the analyzed (phospho-)proteins were not expected. Densitometric analysis of the obtained data was performed using ImageJ software (ImageJ, version 1.52a). For immunoprecipitation, protein adjusted whole-cell lysates were incubated with the indicated antibody overnight at 4°C under continuous agitation. Then, 25 μl packed Protein G-Sepharose beads (GE Healthcare, #GE17-0618-01) were added for 2 h at 4°C under continuous agitation. Beads were pelleted (500 x g, 5 min) and supernatants were removed. The beads were washed in diluted (1/5) lysis buffer thoroughly and the precipitates were separated by SDS-PAGE and analyzed by immunoblotting as described above.

The following antibodies were used:

PI3K p85 (CST, #42925); HSP90 (CST, #4877); p-PLCγ1 (Tyr783) (CST, #14008); p-PKB (Ser473) (CST, #4051); p-JNK (Thr183/Tyr185) (CST, #4668); p-MEK1/2 (Thr217,Ser221) (CST, #2338); p-ERK1/2 (Thr202/Tyr204) (CST #4370); p-Tyr100 (CST, #9411) p-IκBα (Ser32) (CST, #2859); IκBα (CST; #9242) p-c-Kit (Tyr719) (CST, #3391), all purchased from Cell Signaling Technology. GAPDH (#sc-32233); LYN (#sc-7274); SHIP1 (P1C1, #sc-8425); SHIP1 (N1, #sc-6244); GST (#sc-138) were purchased from Santa Cruz Biotechnology.

Supernatant of 4G10 hybridoma cells was used for detection of phosphorylated tyrosines. HRP-coupled secondary antibodies against goat (#P044901), mouse (#P0447) and rabbit (#P0448) were from Agilent (Dako).

### MicroScale Thermophoresis (MST)

Protein labelling for MST: The human active protein kinases LYN (ProQinase, #0358-0000-1) and FYN (ProQinase, #0352-0000-1) were labeled using Monolith™ Protein Labeling Kit RED-TRIS NTA 2^nd^ gen (NanoTemper Technologies, # MO-L018) according to the manufacturer’s instructions.

MicroScale Thermophoresis (MST) binding experiments were carried out with 25 nM Red Tris NTA-labeled FYN in binding buffer (20 mM HEPES pH 8, 100 mM NaCl, 0.05% Tween-20, 1% Glycerol, 2% DMSO) with a range of concentrations of KIRA6 (200 µM to 6.1 nM) at 40% MST (medium) power, 20% LED power in premium capillaries on a Monolith NT.115 pico device at 25°C (NanoTemper Technologies). Red Tris NTA labelled LYN was studied at 25 nM in binding buffer (20 mM HEPES pH 8, 100 mM NaCl, 0.05% Tween-20, 1% Glycerol, 1% DMSO) with a range of concentrations of KIRA6 (100 µM to 3.05 nM) at 60% MST (high) power, 20% LED power in premium capillaries. Data were analyzed using MO.Affinity Analysis software (version v2.3, NanoTemper Technologies) at the standard MST-on time of 30 s. Data fits possessing amplitudes >5 units combined with Signal to Noise levels >5 units were defined as binding events. Aggregation was not observed in the experiments. In order to calculate fraction bound, the ΔFnorm value of each point is divided by the amplitude of the fitted curve, resulting in values from 0 to 1 (0 = unbound, 1 = bound), and processed using the Kaleidagraph software. Error bars represent the standard deviation of 2 independent experiments in (two technical repeats each).

### Nano-differential scanning fluorimetry (nanoDSF)

For modification free thermal unfolding experiments, FYN or LYN were studied at 800 nM in binding buffer (20 mM HEPES pH 8, 100 mM NaCl, 0.05% Tween-20, 1% Glycerol, 1% DMSO) in absence and presence of KIRA6 (100 µM). Samples were pre-incubated in high sensitivity capillaries for 30 min before measurement in the Prometheus NT.48 instrument (NanoTemper Technologies) with 50% sensitivity and a temperature ramp of 1°C / min from 20 to 95°C. Two independent experiments were performed and merged for analysis at 330 nm wavelength using the software PR. StabilityAnalysis (version v1.1, NanoTemper Technologies).

### *In vitro* kinase assay

The LYN kinase assay was performed according to the previously described protocol by Laborlette et al. (Larbolette et al., 1999). Briefly, recombinant GST-fusion protein SH3-P7 was used as a LYN substate. GST-SH3P7 was recombinantly expressed and purified as described by Brummer and colleagues (Brummer et al., 2004). LYN was immunoprecipitated from unstimulated BMMCs with anti-LYN antibody as previously described (Miranda et al., 2016)

The kinase reaction was performed in the presence of 200 mM ATP with 5 µg SH3-P7 and LYN precipitated from 3 x 10^6^ cells for 30 min at RT in 20 µl of kinase buffer containing 40 mM HEPES, pH7.4; 10mM MgCl_2_, 3mM MnCl_2_. DMSO (solvent control) or KIRA6 was pre-incubated 10 min at RT before the addition of ATP. The reaction was terminated by adding 2x Laemmli buffer and subsequently samples were boiled at 95°C for 5 min before being subjected to SDS-PAGE and Western blotting. Phosphorylated SH3-P7 was detected by immunostaining using 4G10 antibody.

### Molecular Modeling techniques

The active states of the three kinases were taken from the Protein Data Bank (Berman et al., 2000) (PDB codes: 1PKG, 2DQ7 and 5XY1 for KIT, FYN and LYN, respectively). The inactive state of KIT was available on the PDB (PDB code 6MOB), while no experimentally determined structure of the inactive (DFG-out) states was available for the human LYN and FYN proteins. Therefore, we generated these using the MODELLER software (Eswar et al., 2008).

LYN(DFG-out) homology model: the structure of LYN bound to an inhibitor (PDB_ID: 5XY1) (Williams et al., 2009) was used as a template with the activation loop deleted (Residues: 383-404). The structure of LCK in complex with type 2 inhibitor imatinib was used in the template to model the DFG-out conformation (PDB_ID 2PL0). We generated 100 models, of which the best-scored (DOPE score) (Shen & Sali, 2006) structure was used in the following steps.

FYN(DFG-out) homology model: we used the active state crystal structure of FYN (PDB_ID: 2DQ7) (Jacobs et al., 2008) as template for the model. The procedure for the activation loop mimics the one described above for LYN.

Docking of KIRA6 to LYN/FYN/KIT: we docked KIRA6 to the respective proteins with Schrödinger’s Glide software (Friesner et al., 2004)(Halgren et al., 2004) in its Induced Fit mode with standard parameters. We chose the ligand centroids of the homology models and the ligands co-crystallized as the origins for the 10x10x10 Å inner bounding box.

Molecular Dynamics Simulations: 200ns of molecular dynamics trajectories using GROMACS (Berendsen et al., 1995) with the AMBER ff99SB-ILDN(Lindorff-Larsen et al., 2010) forcefield were generated. The clustering of the ligand poses was done using the ttclust (Tubiana et al., 2018) python package. The standard parameters for clustering were used. Alignment was performed on the backbone atoms and the ligand atoms were used for the RMSD calculation. The cluster distance metric was performed using the “ward” metric. The ‘autocluster’ parameter -y was set to yes, which uses the elbow criterion for cluster number estimation.

*MM-GBSA.* The estimation of the binding free energies in between KIRA6 binding to the different proteins were performed using the gmx_MMPBSA (Miller et al., 2012; Valdés-Tresanco et al., 2021) tool. We removed the PBC conditions for this and used every frame of the last 100 ns after convergence with the parameters igb=2 and saltcon=0.15. Additionally, we used the interaction entropy approximation implemented in the tool for estimation of the conformational entropy. As a comparison, we also calculated the energetics of the cluster representatives with the single point MMGBSA function of the Schrödinger tool suit.

### Preparation and use of precision-cut lung slices

Precision-cut lung slices (PCLS) were prepared from 8-week-old Wistar rats (220 ± 20 g) obtained from Janvier and kept under controlled conditions (22°C, 55% humidity and 12-h day/night rhythm). Animal experiments were approved by the local ethics committee. Rat PCLS were prepared as previously described (Wohlsen et al., 2001). Rats were sacrificed by an over-dose of pentobarbital i.p. (600 mg/kg). Isolated lungs were filled with pre-warmed agarose solution (0.75%) via the trachea and subsequently chilled with ice. Then lobes were separated and cut into 5 to 10 mm thick tissue segments from which cores were made along the airways, and then cut into 250 ± 20 μm thick slices (Alabama Research and Development). For studies with ovalbumin, the lung slices were incubated overnight with cell culture medium containing 1% of serum from actively sensitized rats, as previously shown (Wohlsen et al., 2001). After overnight culturing, the airways in PCLS were imaged and digitized using a digital video camera. A control picture was taken before addition of DMSO or KIRA6 and after addition of ovalbumin (10 μg/ml) frames were recorded every 30 s for 15 min. The images were analyzed by the image analysis program Optimas 6.5 (Optimas).

### Statistics

Data were generated from at least n = 3 independent experiments. *p*-values were calculated as indicated in the respective figure legends using GraphPad Prism. Data are shown as mean ± SD of n ≥ 3 independent experiments (with n indicated in the respective figure legends). *p*-values of * < 0.05, ** < 0.01, *** < 0.001 and **** < 0.0001 were considered statistically significant. Values higher than a *p*-value of 0.05 were regarded as not significant (ns).

## Results

### KIRA6 represses early and sustained pro-inflammatory functions in Ag-triggered mast cells

The IRE1α/XBP1 arm of the UPR not only secures the increased need for protein folding, but also promotes *Il6* transcription by XBP1s binding to the *Il6* promoter in endotoxin-activated macrophages (Martinon et al., 2010). We sought to investigate if this mechanism holds true for MCs, another important innate immune cell, and particularly focused on *Ship1*-deficient BMMCs. These cells are hyperactive and produce high amounts of pro-inflammatory cytokines in response to FcεRI crosslinking (Fehrenbach et al., 2009). A dependence of IL-6 production should be investigated using the IRE1α inhibitor KIRA6 (Ghosh et al., 2014). Indeed, preliminary experiments showed that a pretreatment with KIRA6 causes attenuation of Ag-triggered IL-6 production/secretion (data not shown). As a matter of routine, we used the same cellular supernatants to also measure for the presence of the lysosomal enzyme β-hexosaminidase, which, in contrast to IL-6, is released immediately after FcεRI activation. Unexpectedly however, severe reduction of secreted β-hexosaminidase was found in the cellular supernatant of KIRA6-pretreated *Ship1*^-/-^ BMMCs (data not shown). Kinetically/mechanistically, no obvious reason could explain why this rapid β-hexosaminidase-releasing degranulation process should depend on the activity of the ER stress sensor IRE1α or the transcription factor XBP1s. Hence, we set out to analyze whether KIRA6 has affects MC activation in an off-target manner.

We pretreated *Ship1*^+/+^ and *Ship1*^-/-^ BMMCs with DMSO (vehicle) or rising concentrations of KIRA6 (0.03 – 1 µM), stimulated with Ag (DNP-HSA), and determined the release of β-hexosaminidase. In agreement with previous data, release of β-hexosaminidase was considerably stronger from *Ship1*^-/-^ compared to *Ship1*^+/+^ BMMCs (Huber et al., 1998) (Gimborn et al., 2005). Intriguingly, KIRA6 inhibited Ag-triggered degranulation in *Ship1*^+/+^ BMMCs starting at a concentration of 0.03 µM, whereas in *Ship1*^-/-^ BMMCs an approximately 30x higher concentration of KIRA6 (1 µM) was necessary to obtain a comparable level of inhibition (Fig. 1A).

**Figure 1:**
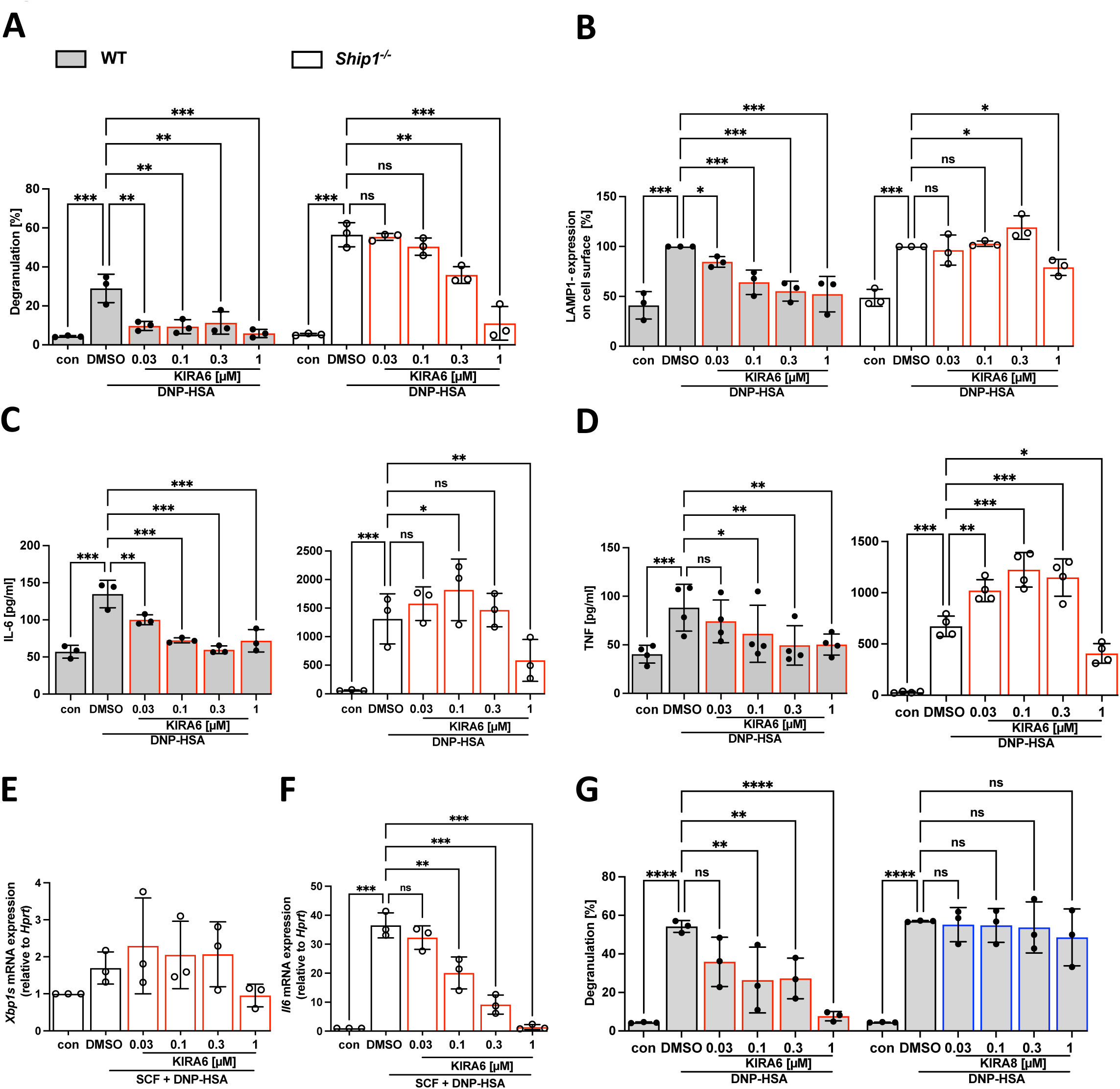
KIRA6 inhibits degranulation and cytokine production of WT and *Ship1*^-/-^ BMMCs. IgE pre-loaded WT (gray bars) or *Ship1^-/-^* BMMCs (white bars) were incubated with solvent control DMSO or the indicated KIRA6 (red line) or KIRA8 (blue line) concentrations for 1 h followed by Ag-stimulation with 20 ng/ml DNP-HSA (A-D, G) or combined stimulation with 100 ng/ml SCF (E, F).(**A**) Degranulation was determined by calculating the enzymatic activity of released β-hexosaminidase in relation to total β-hexosaminidase activity after stimulation of WT and *Ship1^-/-^* cells with DNP-HSA for 30 min (n=3). (**B**) Cell surface localized LAMP1 was detected by flow cytometry in unstimulated and DNP-HSA-stimulated (5 min) WT and *Ship1^-/-^* BMMCs (n=3). (**C**) Secreted IL-6 (n=3) and (**D**) TNF (n=4) was measured by ELISA assays after 4 h of DNP-HSA stimulation from supernatants of stimulated and unstimulated WT and *Ship1^-/-^* cells. (**E**) *Xbp1s* and (**F**) *Il6* mRNA expression was detected by RT-qPCR after combined stimulation with 100 ng/ml SCF 5 min prior the addition of DNP-HSA for 4 h in *Ship1^-/-^*BMMCs (both n=3). *Hprt* served as housekeeping gene to which mRNA expression of genes of interest was normalized to. (**G**) WT BMMCs were pre-incubated with KIRA6 or KIRA8 followed by DNP-HSA stimulation for 30 min and the percentage of released β-hexosaminidase was calculated (n=3). Data are shown as mean +/- SD. Symbols indicate individual values of biological replicates. (**A**), (**B**) and (**G**) two-way ANOVA followed by Sidak’s multiple comparisons test. (**C**), (**D**), (**E**) and (**F**) RM one-way ANOVA followed Dunnett test to correct for multiple comparison. * p<0.05, ** p<0.01, *** p<0.001, **** p<0.0001, ns indicates non significance.

To support our findings, we additionally made use of the LAMP1 assay for interrogating Ag-triggered secretion. LAMP1, a.k.a. CD107a, is expressed in the membrane of secretory lysosomes and is thus extracellularly not detectable in resting MCs by fluorescently labeled anti-LAMP1 antibodies. Upon Ag stimulation, vesicle membranes fuse with the plasma membrane and hence allow for detection of LAMP1 using flow cytometry. As shown in Fig. 1B, results comparable to the β-hexosaminidase assay were obtained in *Ship1*^+/+^ and *Ship1*^-/-^ BMMCs. *Ship1*^+/+^ BMMCs once again were considerably more sensitive to KIRA6 pretreatment compared to *Ship1*^-/-^ cells.

Next, we investigated the effect of KIRA6 on Ag-induced pro-inflammatory cytokine production in *Ship1*^+/+^ and *Ship1*^-/-^ BMMCs by measuring secreted IL-6 and TNF. As expected, *Ship1*^-/-^ BMMCs upon Ag stimulation produced dramatically more cytokines than *Ship1*^+/+^ BMMCs (Kalesnikoff et al., 2002)(Fehrenbach et al., 2009) (Fig. 1C&D). Again, KIRA6-mediated inhibition was more efficient in *Ship1*^+/+^ compared to *Ship1*^-/-^ BMMCs (Fig. 1C&D). Intriguingly, in *Ship1*^-/-^ BMMCs low concentrations of KIRA6 first resulted in an increase in cytokine production before a significant reduction occurred with 1 µM KIRA6 (Fig. 1C&D). As mentioned above, the immediate response of degranulation kinetically does not coincide with an ER stress-induced activation of the UPR sensor IRE1α, hence ruling out KIRA6-mediated inhibition of IRE1α as molecular reason for suppression of degranulation. The process of cytokine production, however, could fit to activation of IRE1α and resulting *Xbp1* splicing, as it has been demonstrated in LPS-stimulated macrophages (Martinon et al., 2010). Therefore, we measured *Xbp1* splicing in Ag- and stem cell factor (SCF)-triggered *Ship1*^-/-^ BMMCs and the potential inhibitory effect of KIRA6. As shown in Fig. 1E, virtually no induction of *Xbp1* splicing and thus, no effect of KIRA6 was measurable. However, successful Ag and SCF co-stimulation and inhibition by KIRA6 was verified by measuring production of *Il6* mRNA (Fig. 1F). Importantly, KIRA6 was only able to reduce degranulation in MCs stimulated with Ag alone or in combination with SCF (Suppl. Fig. 1A). To finally rule out participation of IRE1α in Ag-triggered degranulation, we compared the effect of KIRA6 pretreatment with the efficiency of the more recent and selective KIRA8 (Morita et al., 2017)). Indeed, compared to KIRA6, KIRA8 did not suppress β-hexosaminidase release even at 1 µM (Fig. 1G). Nanomolar efficacy of KIRA8 was verified in HMC-1.2 MC leukemia cells stressed with tunicamycin causing strong *XBP1* splicing (Suppl. Fig. 1B). In conclusion, the IRE1α inhibitor KIRA6 suppresses immediate and sustained Ag-triggered MC activation in an IRE1α-independent manner.

### Early Ag-triggered signaling processes are sensitive to KIRA6

Degranulation of MCs can also be triggered by a combination of the Ca^2+^ ionophore A23187 and of the PKC activator PMA (Sagi-Eisenberg et al., 1985). Comparison of the KIRA6 effect on degranulation of BMMCs in response to Ag versus the combination of A23187+PMA was supposed to allow to determine if the target of KIRA6 is localized up-or downstream of the Ca^2+^ mobilization step mandatory for degranulation. In contrast to Ag, A23187 + PMA-induced degranulation of WT BMMCs was not suppressed by KIRA6 (Fig. 2A). This suggested that KIRA6 inhibits an immediate step in FcεRI-induced signal transduction, which is preceding Ca^2+^ mobilization and/or PKC activation. This might be activation of PLCγ, which hydrolyzes PI(4,5)P_2_ to yield IP_3_ and DAG responsible for Ca^2+^ release from the ER and PKC activation at the plasma membrane, respectively. Confirming this assumption, KIRA6 dose-dependently and efficiently suppressed the activating phosphorylation of PLCγ1 at Tyr783 in response to Ag stimulation (Fig. 2B). In agreement, KIRA6 repressed Ag-triggered Ca^2+^ mobilization in a dose-dependent fashion (Fig. 2C; Suppl. Fig. 2). In conclusion, KIRA6, put on the market as an IRE1α inhibitor, is capable of effectively suppressing Ag-triggered MC signaling and effector responses apparently by inhibiting an additional target protein (e.g. an FcεR1an FcεR1-associated kinase).

**Figure 2:**
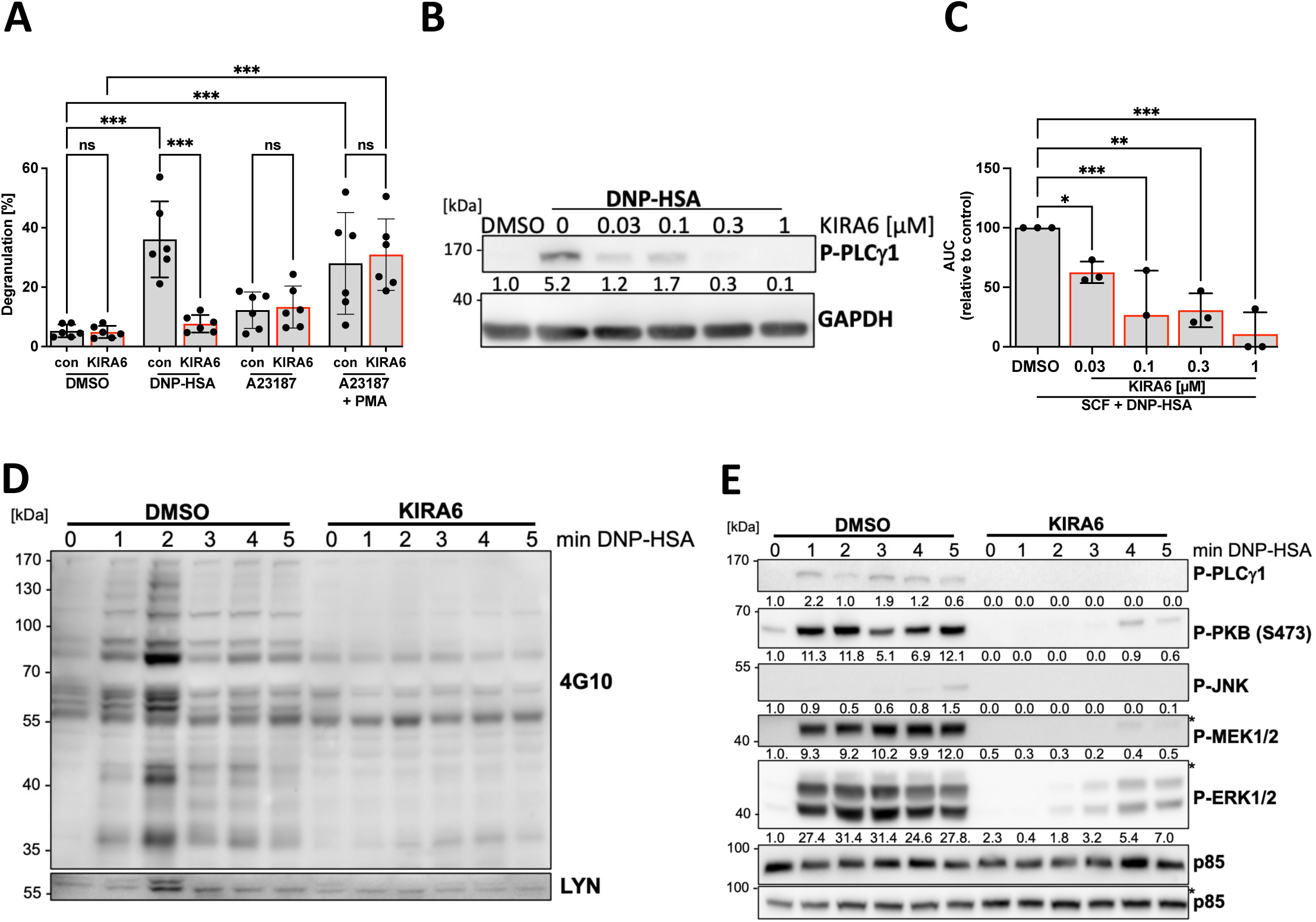
KIRA6 inhibits FcεRI-dependent signaling in WT BMMCs. IgE pre-loaded WT BMMC (gray bars) were incubated with the solvent control DMSO or the indicated KIRA6 (red line) concentrations for 1 h followed by incubation with the indicated stimuli. (**A**) Degranulation was induced by DNP-HSA (20 ng/ml), A23187 (100 ng/ml) or the combination of A23187 (100 ng/ml) and PMA (100 ng/ml) for 30 min and the percentage of released β-hexosaminidase in relation to total β-hexosaminidase was calculated (n=6).(**B**) IgE pre-loaded WT BMMCs were starved overnight, incubated with indicated KIRA6 concentrations or DMSO for 1 h and subsequently stimulated with DNP-HSA (20 ng/ml) for 1 min. PLCγ1 phosphorylation (pTyr783) was assessed by Western blot using a phospho-specific antibody. GAPDH served as loading control (n=3). (**C**) Ca^2+^ mobilization was induced by SCF and DNP-HSA co-stimulation and determined by the ratio of the Ca^2+^-bound/Ca^2+^-unbound using the Ca^2+^-sensitive fluorophores Fluo-3 and FuraRed. The graph shows the normalized comparison of the area under the curve (AUC) values representing total Ca^2+^ mobilized within 4 min after stimulation (n=3). (**D**) Representative Western blot showing global tyrosine phosphorylation after 20 ng/ml DNP-HSA stimulation at the indicated time points using 4G10 antibodies. LYN was detected on the same membrane using an anti-LYN antibody. LYN expression served as loading control (n=3). (**E**) Representative Western blot showing DNP-HSA-induced phosphorylation of PLCγ1, PKB, JNK, MEK1/2 and ERK1/2 detected by Western blot using phospho-specific antibodies. p85 served as a loading control (n=3). (**B**) and (**E**) Numbers indicate mean values obtained from densitometry analyses. Data are shown as mean +/- SD. Symbols indicate individual values of biological replicates. (A) Two-way ANOVA followed by Sidak multiple comparisons test. (**B, C**) RM one-way ANOVA followed by Dunnett’s test to correct for multiple comparison. * p<0.05, ** p<0.01, *** p<0.001, ns indicates non significance.

In previous work, we found that *Lyn*-deficiency results in a comparable reduction of PLCγ phosphorylation and Ca^2+^ mobilization as currently observed for KIRA6 action (Miranda et al., 2016)). Indeed, analysis of total substrate Tyr phosphorylation revealed almost its complete inhibition by KIRA6 pretreatment in *Ship1*^+/+^ and *Ship1*^-/-^ BMMCs already as early as 1 min after addition of Ag (Fig. 2D; Suppl. Fig. 3A), again hinting at LYN as a KIRA6 target. A further characteristic of Ag-stimulated *Lyn*^-/-^ BMMCs was the delayed and weakened phosphorylation of ERK1/2 (Miranda et al., 2016), which was also found in KIRA6-treated *Ship1*^+/+^ and *Ship1*^-/-^ BMMCs, in addition to a repression of MEK1/2, JNK, and PKB phosphorylation (Fig. 2E; Suppl. Fig. 3B). In conclusion, KIRA6 suppresses immediate early Ag-triggered signaling processes, a pattern which is suggestive for the SFK LYN being the/an additional target of KIRA6.

### Pharmacological inhibition of LYN by KIRA6

A specific substrate of LYN in Ag-triggered BMMCs is the lipid phosphatase SHIP1, which attenuates PI3K signaling and IgE-mediated effector functions (Miranda et al., 2016)(Hernandez-Hansen et al., 2004). In accordance with LYN being a potential target of KIRA6, pretreatment of BMMCs with KIRA6 resulted in complete abrogation of basal and Ag-induced Tyr phosphorylation of SHIP1 (Fig. 3A).

**Figure 3:**
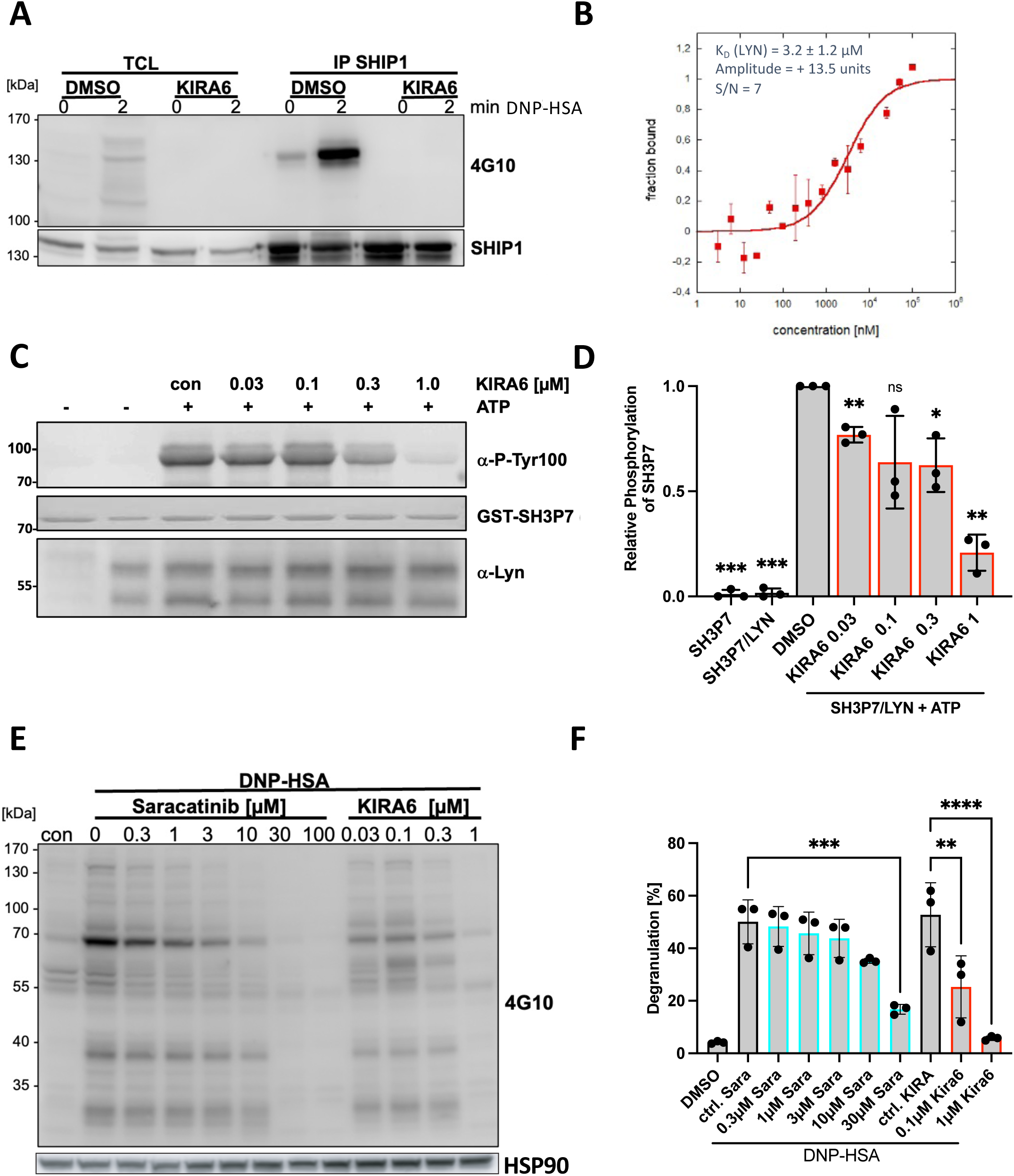
KIRA6 efficiently inhibits LYN kinase activity. **(A)** IgE-sensitized WT BMMCs were pre-incubated with 1 µM KIRA6 for 60 min before cells were left unstimulated or stimulated with 20 ng/ml DNP-HSA for 2 min. SHIP1 was immunoprecipitated from cell lysates and tyrosine phosphorylation of SHIP1 was detected on WB using 4G10 antibodies. A comparable amount of SHIP1 in the TCL and in the precipitates was confirmed by staining with anti-SHIP1 antibody (n=3). (B) Concentration dependent ligand binding of KIRA6 to LYN was evaluated with MicroScale Thermophoresis (MST) to calculate the K_D_. (C) LYN kinase activity was determined *in vitro* by the detection of the phosphorylated LYN substrate GST-SH3P7 using a pTyr100-sepcific antibody. The substrate was incubated with precipitated LYN and the reaction was performed in the presence of 200 mM ATP for 30 min at RT. KIRA6 or the solvent DMSO were pre-incubated for 10 min before the addition of ATP. Ponceau S was used to stain recombinant GST-SH3P7 on the WB membrane and an anti-LYN antibody to show comparable amounts of LYN in each sample. (**D**) Relative SH3P7 phosphorylation of independently performed LYN kinase assays was quantified in comparison to the positive control (DMSO) using the solvent DMSO with LYN and SH3P7 in the presence of ATP. Relative phosphorylation intensity was determined by densitometry analysis (n=3). (**E**) and (**F**) IgE-sensitized WT BMMCs were incubated with indicated concentrations of Saracatinib and KIRA6 for 60 min before DNP-HSA stimulation. (**E**) Global tyrosine phosphorylation following DNP-HSA stimulation and KIRA6 or Saracatinib pretreatment (n=3). (**F**) The influence of Saracatinib (Sara) and KIRA6 on DNP-HSA-induced degranulation was calculated by the percentage of released in relation to total β-hexosaminidase (n=3). (**D**) and **(F)** Data are shown as mean +/- SD. Symbols indicate individual values of biological replicates. **(D)** One sample t-test in comparison to the positive control (DMSO, LYN, SH3P7, ATP). (**F**) was analyzed by ordinary one-way ANOVA followed by Sidak’s multiple comparison test. * p<0.05, ** p<0.01, *** p<0.001, **** p<0.0001, ns indicates non significance.

We determined whether KIRA6 can directly bind to recombinant LYN *in vitro* and calculated the *K_d_* of this interaction. Performing MST using fluorescently labeled LYN (Gorny et al., 2022), the calculated *K_d_* of KIRA6 was 3.2 ± 1.2 µM (Fig. 3B). Moreover, interaction between LYN and KIRA6 was verified by means of nanoDSF (a label-free thermal-shift assay technique) to study protein denaturation as function of temperature (Gorny et al., 2022), which proved a distinct change in thermal unfolding of recombinant LYN (ΔIP1 = -3.1°C) in the presence of KIRA6 (Suppl. Fig. 4A). Following, we performed an in vitro kinase assay using immunoprecipitated LYN from BMMC lysates and a GST fusion protein containing the full-length LYN substrate SH3P7 as substrate (Larbolette et al., 1999) We could clearly show that KIRA6 inhibits LYN with an *IC_50_*of approximately 300 nM (Fig. 3C&D; Suppl. Fig. 4B), illustrating that measurement of direct binding of KIRA6 to recombinant LYN protein using MST correlates qualitatively with *in vitro* analysis (IVKA).

Being now confident that KIRA6 is a LYN inhibitor in the sub-micromolar range, we compared its efficiency with the potent SFK inhibitor, Saracatinib (AZD-0530), which has been shown in cell-free assays to inhibit LYN with an *IC*_50_ of 5 nm (Green et al., 2009). However, in different cellular assays, Saracatinib was used at concentrations between 1 and 10 µM (Yamaki et al., 2020)(Chang et al., 2008)(Chua et al., 2015). Analysis of Ag-triggered substrate Tyr phosphorylation by Western blotting as well as measurement of β-hexosaminidase release revealed a considerable difference in effectiveness between KIRA6 and Saracatinib, with KIRA6 being more efficient by a factor of approximately 30 (Fig. 3E&F). In conclusion, our findings indicate that KIRA6 binds to and inhibits the SFK LYN in the sub-micromolar range, effectively suppressing Ag-triggered MC activation.

### Use of *Lyn*-deficient BMMCs corroborates LYN inhibition by KIRA6 and suggests additional suppression of FYN at higher concentrations

Given that KIRA6 is a potent LYN inhibitor, it should not suppress Ag-induced degranulation or cytokine production in LYN-deficient MCs, at least not at low nanomolar concentrations effective in WT cells. Therefore, we generated *Lyn*^-/-^ BMMCs and compared their Ag-triggered effector and signaling functions to WT (*Lyn*^+/+^) BMMCs in the presence or absence of KIRA6. First, we analyzed Ag-induced degranulation (β-hexosaminidase release) in the presence of increasing concentrations of KIRA6. While KIRA6 treatment of *Lyn*^+/+^ cells caused progressive attenuation of degranulation between 0.03 and 0.3 µM of KIRA6, no such effect was observed in *Lyn*^-/-^ BMMCs (Fig. 4A). At 1 µM of KIRA6, however, complete suppression of degranulation was observed in both *Lyn*^+/+^ and *Lyn*^-/-^ cells, suggesting that at this concentration one or more additional KIRA6-sensitive kinases, which are positively regulating MC degranulation, are inhibited, with the SFK FYN being a meaningful candidate (Parravicini et al., 2002). We further analyzed the effect of KIRA6 on Ag-triggered production of IL-6 and TNF in *Lyn*^+/+^ and *Lyn*^-/-^ BMMCs. As expected, *Lyn*^-/-^ BMMCs, compared to *Lyn*^+/+^ BMMCs, produced considerably higher amounts of the pro-inflammatory cytokines (Fig. 4B&C) (Miranda et al., 2016). As observed for degranulation, a comparable pattern could be demonstrated with respect to IL-6 (Fig. 4B) and TNF production (Fig. 4C) in KIRA6-pretreated (0.03 – 0.3 µM) *Lyn*^+/+^ and *Lyn*^-/-^ BMMCs. Unexpectedly, secretory responses in LYN-deficient BMMCs were even promoted by KIRA6 in the range of 0.03 – 0.3 µM (Fig. 4A-C), which might depend on differential sensitivity for KIRA6 of the different SFKs expressed in MCs (Suppl. Fig. 6C) as well as respective differences between *Lyn*^+/+^ and *Lyn*^-/-^ BMMCs. Nevertheless, our observations using *Lyn*^-/-^ BMMCs clearly corroborate the LYN-specific action of KIRA6 at sub-micromolar concentrations.

**Figure 4:**
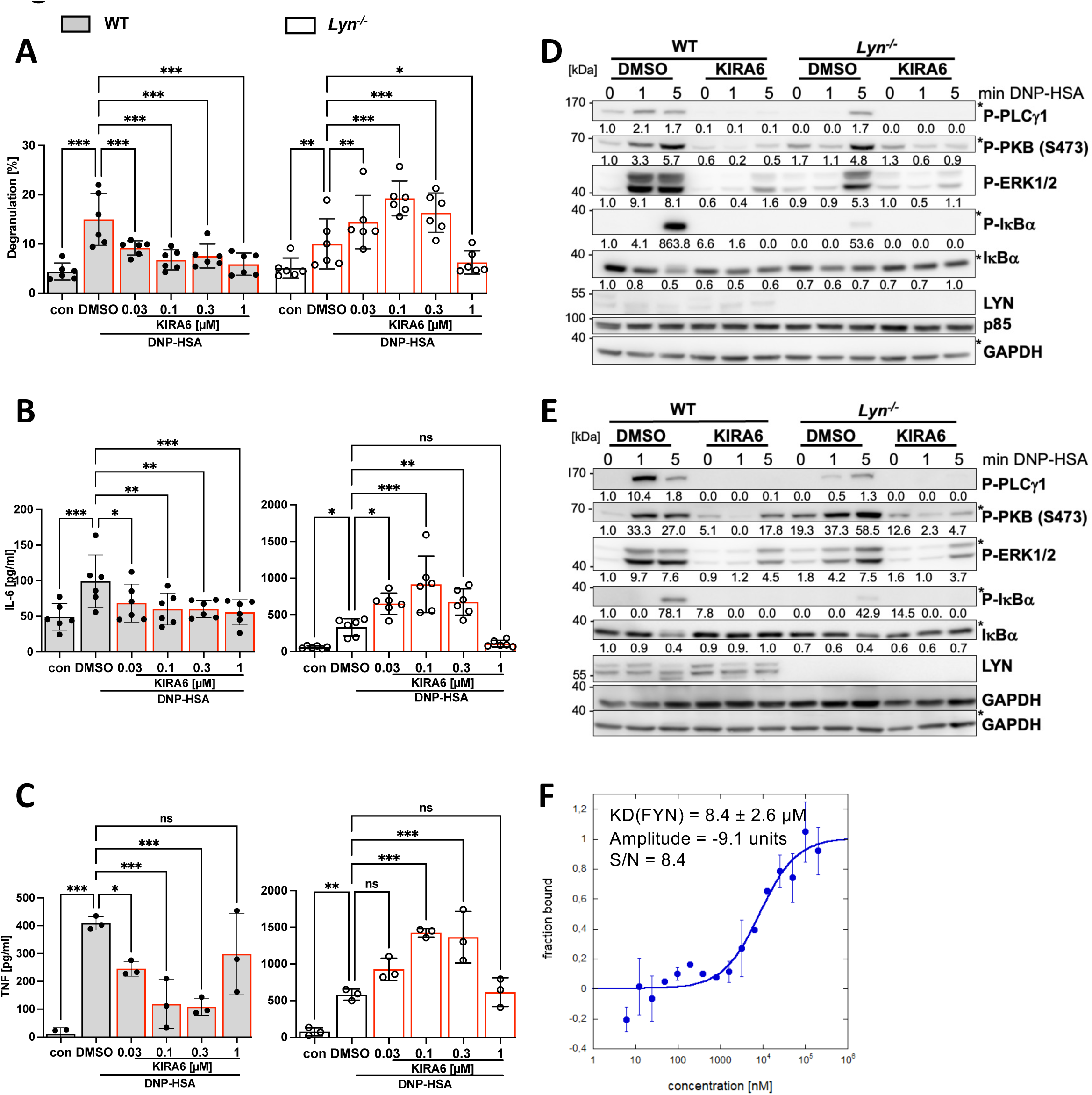
KIRA6 additionally affects FYN activation. IgE pre-loaded WT BMMC (gray bars) or *Lyn^-/-^* BMMCs (white bars) were incubated with the solvent control DMSO or the indicated KIRA6 (red line) concentrations for 1 h followed by antigen stimulation with 20 ng/ml DNP-HSA. (**A**) Degranulation was calculated by the percentage of released in relation to total β-hexosaminidase after stimulation with DNP-HSA for 30 min (n=5). (**B**) Released IL-6 and (**C**) TNF was measured by ELISA assays after KIRA6 pretreatment followed by DNP-HSA stimulation for 4 h (both n=6). (**D**) 20 ng/ml DNP-HSA or (**E**) 200 ng/ml DNP-HSA were used to stimulate WT or Lyn^-/-^ BMMCs for the indicated time points to induce FcεRI-dependent phosphorylation of PLCγ1, PKB, ERK1/2 and IκBα shown on WB using phospho-specific antibodies. P85 and GAPDH served as a loading control. Phosphorylation signals detected on the same membrane are marked with an asterisk. Numbers indicate mean values obtained from densitometry analyses (n=3). (**F**) Concentration-dependent ligand binding of KIRA6 to FYN was evaluated with MicroScale Thermophoresis (MST) to calculate the K_D_. (**A**), (**B**) and (**C**) Data are shown as mean +/- SD. Symbols indicate individual values of biological replicates. (**A**) Two-way ANOVA followed by Dunnett’s multiple comparisons test. **(B)**,), **(C)** RM one-way ANOVA followed by Dunnett’s test to correct for multiple comparison. * p<0.05, ** p<0.01, *** p<0.001, **** p<0.0001, ns indicates non significance.

The further analysis of Ag-triggered signaling events in *Lyn*^+/+^ and *Lyn*^-/-^ BMMCs in the absence or presence of KIRA6 completed the picture (Fig. 4D). Ag-induced phosphorylation of PLCγ1, PKB, ERK1/2, and IκBα were markedly reduced in the absence of LYN (Miranda et al., 2016)), and the remaining phosphorylations in *Lyn*^-/-^ BMMCs were suppressed to background levels by 1 µM KIRA6 (Fig. 4D).

Intriguingly, LYN is capable of both positively and negatively regulating FcεRI-mediated MC activation with higher concentrations of crosslinking stimulus shifting the balance to inhibitory functions (Xiao et al., 2005). Hence, particularly PKB is stronger phosphorylated in *Lyn*^-/-^ BMMCs in response to high Ag concentrations (Xiao et al., 2005)(Miranda et al., 2016). Since PKB phosphorylation is dependent on PI3K, the activation of which has been proven to require FYN activity (Parravicini et al., 2002), we made use of this signaling system to interrogate the potential susceptibility of FYN for inhibition by KIRA6. Increasing the antigen stimulus (DNP-HSA) from 20 ng/ml (Fig. 4D) to 200 ng/ml (Fig. 4E) resulted in the expected enhanced phosphorylation of PKB in *Lyn*^-/-^ compared to *Lyn*^+/+^ BMMCs. This increased, FYN-dependent PKB phosphorylation (Parravicini et al., 2002) could be abrogated by KIRA6 almost completely (Fig. 4E). Moreover, direct binding of KIRA6 to recombinant FYN was analyzed using MST and a *K_d_* of 8.4 ± 2.6 µM was determined (Fig. 4F). Performance of nanoDSF proved interaction between FYN and KIRA6 as well (Suppl. Fig. 4A). Hence, these data strongly suggested that FYN can be inhibited by 1 µM KIRA6. In conclusion, our data reveal that KIRA6 is a potent LYN inhibitor and additionally a reasonable FYN inhibitor, enabling KIRA6 by this combined action to efficiently suppress pro-inflammatory FcεRI-mediated MC activation.

To additionally exclude that the need for higher KIRA6 concentrations to inhibit activation of *Lyn*^-/-^ BMMCs is due to a potential general KIRA6 insensitivity of these cells, we analyzed the effect of KIRA6 on *Lyn*^+/+^ and *Lyn*^-/-^ BMMCs treated with the LYN-independent ER stressor thapsigargin. Thapsigargin inhibits the SERCA located in the ER membrane resulting in an immediate increase of the cytosolic Ca^2+^ concentration, which causes IgE-independent MC activation (Huber et al., 2000). *Lyn*^+/+^ and *Lyn*^-/-^ BMMCs were pretreated with increasing concentrations of KIRA6, subsequently treated with thapsigargin, and production of *Xbp1s* and *Il6* mRNA (Suppl. Fig. 5A-D) as well as IL-6 and TNF protein (Suppl. Fig. 5E&F) were measured. Both cell types produced comparable amounts of *Xbp1s* and *Il6* mRNA as well as IL-6 and TNF protein, and 1 µM KIRA6 was needed for significant reduction of the different MC responses. These data indicate that *Lyn*^+/+^ and *Lyn*^-/-^ BMMCs, in principle, are comparably sensitive to KIRA6 when addressing a LYN-independent signaling process.

### KIRA6 suppresses MC activation in an *ex vivo* model as well as in human MCs

To extend our study to IgE-mediated MC activation in a tissue-type situation, we made use of a model of precision-cut lung slices (PCLS), interrogating the inhibitory effect of KIRA6 on allergen-induced bronchoconstriction in rat lung slices. Due to their intact microanatomy, PCLS allow for the investigation of whole-lung functions in a reproducible manner (Held et al., 1999). PCLS were pretreated with serum from rats sensitized with ovalbumin, and allergen-induced bronchoconstriction was measured in the absence or presence of increasing concentrations of KIRA6. Allergen-induced bronchoconstriction was significantly suppressed by 1 µM KIRA6 (Fig. 5A). The need for a higher drug concentration compared to experiments with BMMCs is most likely due to reduced permeability of the tissue preparation. As control, methacholine-induced bronchoconstriction, which is mediated by the muscarinic acetylcholine receptor, was not affected by preincubation with KIRA6 (Suppl. Fig. 6A).

**Figure 5:**
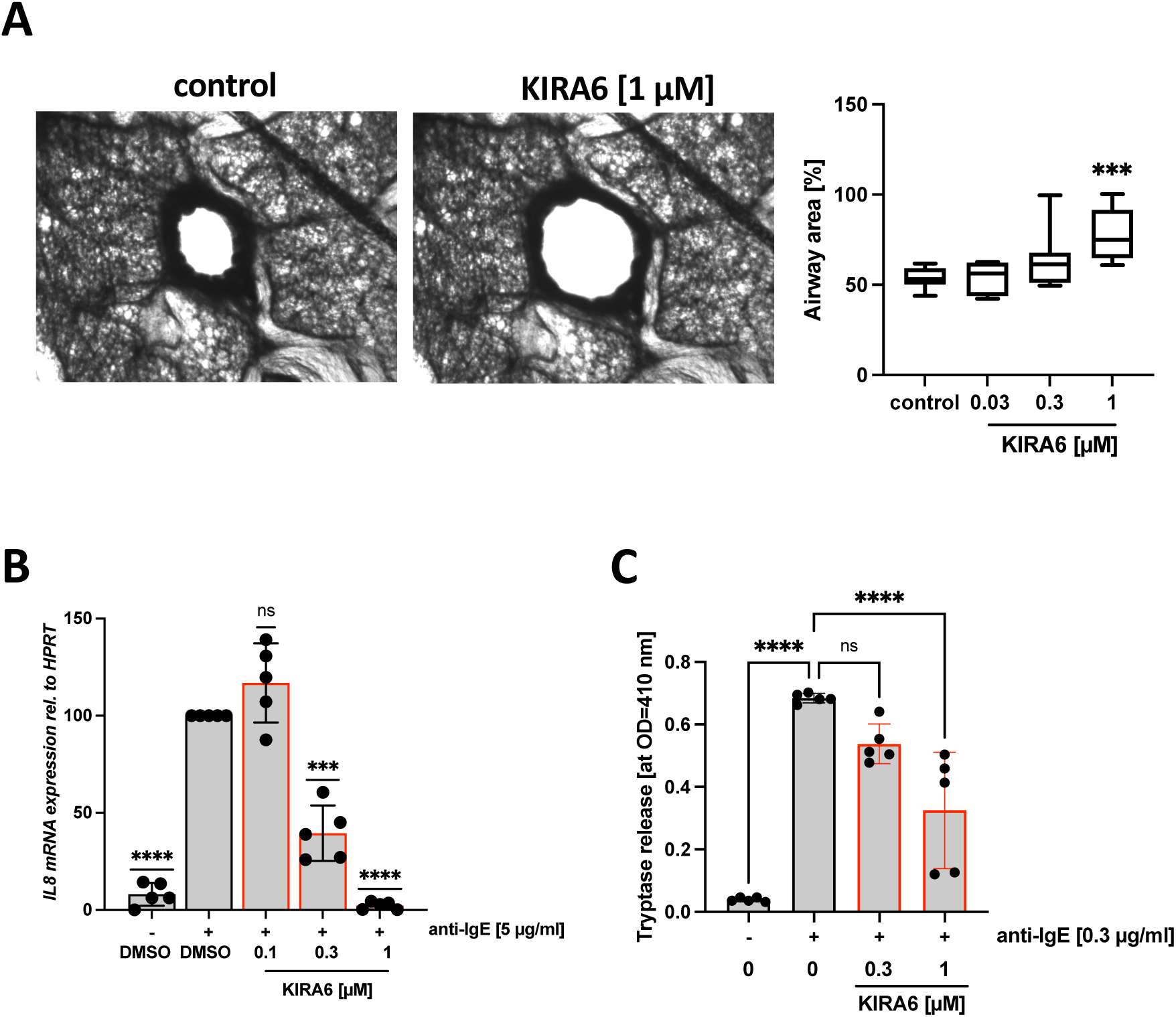
KIRA6 inhibits mast cell function *in vivo* and different *in vitro* model systems. **(A)** Airway contractions to ovalbumin in rat precision-cut lung slices. IgE-sensitized rat precision-cut lung slices were incubated with the solvent control DMSO or the indicated KIRA6 concentration for 5 min prior to challenge with OVA (10 μg/ml) for 15 min and the degree of bronchoconstriction was determined by evaluation of airway area. The images show airway contraction after pre-incubation with the solvent control DMSO (left) or 1 µM KIRA6 (right) and challenge with OVA. The graph shows the response of airway contractions in the presence of the solvent DMSO and the depicted KIRA6 concentration. Contractions are expressed as the decrease of airway area (%) compared to the initial airway area (n=10). (**B**) IL-4- and IgE-primed ROSA ^KIT^ ^WT^ cells were pretreated with DMSO or KIRA6 for 1 h and subsequently stimulated or not with 5 µg/ml α-IgE for 60 min. *IL8* expression was measured by RT-qPCR. *HPRT* served as housekeeping gene to which *IL8* expression was normalized to. (**C**) IgE-sensitized CBMCs were pretreated with DMSO or indicated KIRA6 concentration and stimulated with 5 µg/ml α-IgE for 30 min. Degranulation was evaluated by determining the release of tryptase by an enzymatic assay. (**A**), (**B**) and (**C**) Data are shown as mean +/- SD. Symbols indicate individual values of biological replicates. (**A**), (**B**), (**C)** ordinary one-way ANOVA followed by Dunnett’s test to correct for multiple comparison. * p<0.05, ** p<0.01, *** p<0.001, **** p<0.0001, ns indicates non significance.

Next, we aimed to analyze the inhibitory action of KIRA6 on human MCs. First, we used the SCF-dependent human MC line ROSA^KIT^ ^WT^ as a model system. IgE pre-loaded ROSA^KIT^ ^WT^ MCs were activated by anti-IgE-triggered FcεRI crosslinking in the absence or presence of KIRA6, and *IL8* mRNA expression was measured by RT-qPCR. Significant suppression was already measurable using 0.3 µM KIRA6 (Fig. 5B). Finally, not to only rely on a human MC cell line we utilized IgE-sensitized CBMCs, activated them using anti-IgE antibodies, and measured release of the protease tryptase. Again, KIRA6 (1 µM) was able to completely inhibit the degranulation process (Fig. 5C). In conclusion, KIRA6 treatment was able to inhibit LYN/SFK-dependent MC activation not only in murine BMMCs but also in the *ex vivo* model of bronchoconstriction in PCLS as well as in different human MC models in response to Ag/IgE-mediated activation.

### Molecular modeling reveals preferred KIRA6 interaction with inactive SFKs

Since our research indicated strong interaction of KIRA6 with three tyrosine kinases of prominent importance for MC activation (LYN and FYN [this work], and KIT (Mahameed et al., 2019) as well as their inhibition, we aimed in a final step to provide atomistic level data for these interactions. We modeled the binding of KIRA6 to both an active and an inactive conformation of the three kinases LYN, FYN and KIT. The main factor for discriminating the functional, active state of kinases like FYN, LYN, or KIT from the inactive state has traditionally been the position of a conserved DFG (or, rarely, D[LWY]G) motif in its activation loop (Kufareva & Abagyan, 2008). While most inhibitors bind to the active (DFG-in) state, so-called type-II inhibitors bind to a distinct DFG-out conformation and occupy an additional hydrophobic pocket (Kufareva & Abagyan, 2008)(Vijayan et al., 2015). KIRA6 was docked with on both DFG-in and DFG-out conformations of FYN, LYN, and KIT (please see the methods section for more information on the modeling of these conformations). After performing MD simulations to relax the structure and MM GBSA-calculations (see methods for details) to estimate the binding affinity, it turned out that KIRA6 has a preference to bind the inactive conformation of the three kinases with a slight preference for KIT and LYN kinases in comparison to FYN (see Table 1).

**Table 1:** Energetics of the last 100ns of the converged MD trajectories. ΔH and ΔG value, as well as, Glide SP docking scores of KIRA6 docked to LYN, FYN and KIT are listed.

Specifically, we noticed that while the ligand is nearly fully enclosed in the inactive conformation (Fig. 6B&D), significant parts of the ligand are solved-exposed in the active conformation (see, for example, the Trifluormethyl-phenyl moiety in Fig. 6A&C). The Trifluormethyl-phenyl moiety can make use of the additional pocket that is opened by the DFG-out conformation in the inactive state. In particular, there is an additional hydrogen bond between LYN and KIRA6 in the inactive (ASP385, GLU320, MET322) conformation vs. the active (2x MET322). These factors can explain the difference in docking score between the active (-6.5) and inactive (-14.4) states in KIRA6 binding to LYN.

**Figure 6:**
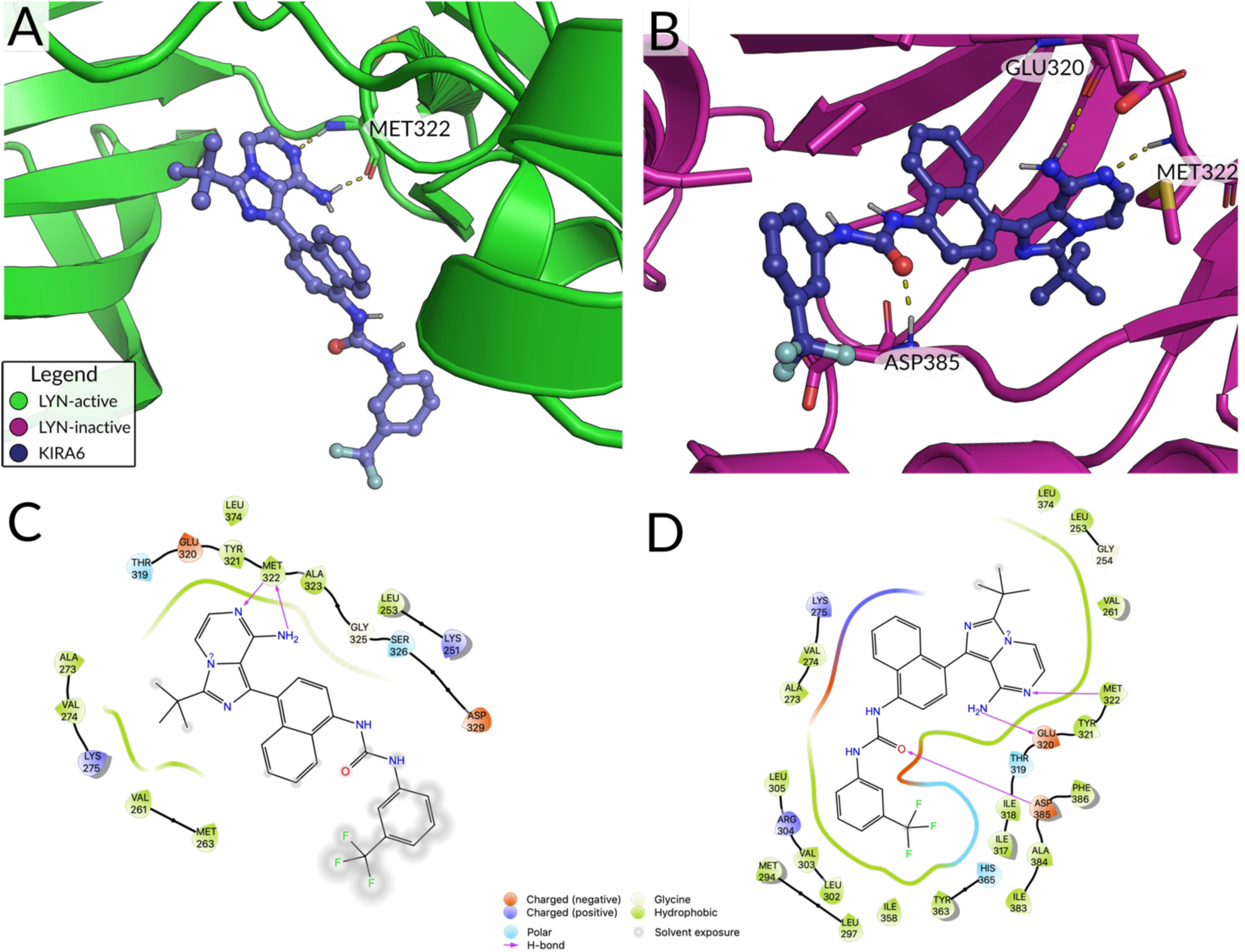
2D and 3D estimated binding poses of KIRA6 binding to the LYN protein. Predicted binding conformation of KIRA6 in active (DFG-in) (**A**) and inactive (DFG-out) (**B**) LYN. Hydrogen bond interactions are visualized with yellow dotted lines. (**C**) and (**D**) 2D representations of the binding poses shown above.

## Discussion

In the present report we demonstrate that the IRE1α inhibitor KIRA6 acts as a very effective inhibitor for the SFK LYN in the nanomolar range, with LYN being one of the main activating and regulatory Tyr kinases in the context of FcRIε signal transduction in MCs. This was demonstrated, amongst others, by the KIRA6-mediated inhibition of LYN-dependent processes including Ag-induced substrate Tyr phosphorylation, Ca^2+^ mobilization, and degranulation. These effects are independent of IRE1α, since i) the concentration required for effective inhibition of IRE1α is higher than the concentration needed for inhibition of LYN-dependent processes, ii) the more effective and selective IRE1α inhibitor KIRA8 (Morita et al., 2017) did not suppress LYN-dependent MC degranulation at a concentration 10-times higher than needed for IRE1α inhibition, and iii) kinetics of such immediate Ag-triggered processes are, in principle, too fast for a molecular involvement of IRE1α. Furthermore, MST and nanoDSF measurements, as well as IVKAs convincingly showed that KIRA6 is structurally and functionally interacting with LYN. Moreover, and consistent with the functional data, a molecular modeling approach verified the close interaction of KIRA6 with the catalytic center of LYN in its inactive state. The LYN-dependence of the KIRA6-mediated effects was further confirmed using *Lyn*^-/-^ BMMCs, where KIRA6, at concentrations effective in WT BMMCs, did not suppress Ag-induced degranulation and cytokine production.

Whereas Ag-triggered effector functions in WT MCs were diminished by KIRA6 in a dose-dependent manner, suppression was only evident at a high concentration of 1 µM in *Lyn*^-/-^ BMMCs, most likely reflecting inhibition of FYN. LYN appears to have a dominant role in the family of SFKs, since it regulates the phosphorylation by the kinase CSK of the C-terminal Tyr residue of other SFKs, and thus their attenuation. Concurring with the importance of LYN and CSK for MC signaling and regulation of SFKs, our Next Generation Sequencing analysis of BMMCs corroborated dominant expression of *Lyn* and *Csk* over *Fyn*, *Hck*, and *Src* (Suppl. Fig. 7B). Odom et al. have shown that Tyr phosphorylation of the membrane-spanning CSK-binding protein (CBP; a.k.a. PAG85) is drastically reduced in *Lyn^-^*^/-^ compared to WT BMMCs, resulting in reduced interaction between CBP and CSK, entailing weaker C-terminal Tyr phosphorylation of FYN and thus its enhanced activity (Odom et al., 2004). In agreement, the SFKs LYN, FYN, HCK, and SRC have been demonstrated to associate with CBP (Kawabuchi et al., 2000). Since our MST, nanoDSF and molecular modeling approaches pointed at KIRA6 being a more efficient inhibitor of LYN than of FYN, the need for the higher concentration of KIRA6 to suppress FcεRI-mediated degranulation and cytokine production in *Lyn*^-/-^ BMMCs goes with such mechanism. This CBP-CSK-driven negative regulation of SFKs was initially described in T-lymphocytes (Brdiclka et al., 2000) and was mostly accepted to take place in further immune cells. Interestingly however, thoroughly analyzing CBP knockout and knockdown BMMCs, Draberova et al. identified a positive function for CBP as well with respect to Ag-triggered effector mechanisms (Draberova et al., 2014), suggesting a more complex regulation of SFK-mediated MC activation. In this regard, the affinity of LYŃs SH2-domain for a phospho-Tyr residue within CBP was found to be distinctly higher than for its phosphorylated C-terminal Tyr508 (Ingley, 2008), indicating that CSK-mediated phosphorylation of LYN (and maybe of other SFKs as well) does not automatically result in inactivation of LYN.

Our initial observation that KIRA6 inhibits Ag-triggered MC activation was made in *Ship1*^-/-^ BMMCs, where, compared to wildtype BMMCs, a higher concentration of KIRA6 was necessary for significant suppression of MC effector functions. *Ship1*^-/-^ BMMCs stand out due to their dramatic accumulation of the PI3K-generated signaling phospholipid, PIP_3_, and consecutive recruitment/activation of PIP_3_-dependent proteins, particularly PKB (Scheid et al., 2002) (Huber et al., 1999), via their PH-domains. Interestingly, FcRεI-induced PI3K activation demands the functional interaction between the SFK FYN, PI3K, and the adaptor protein GAB2 (Parravicini et al., 2002); the latter translocates to the plasma membrane using its PIP_3_-binding PH-domain, potentially organizing an activation-stabilizing/prolonging feed-forward circuit. Hence, SHIP1 deficiency could promote the GAB2-FYN axis and render the cells less dependent on LYN activity. Thus, as observed in our study *Ship1*^-/-^ BMMCs, would be less susceptible to inhibition by low concentrations of KIRA6. In this regard, differences in the LYN/FYN ratio between types of (mast) cells could define susceptibility to inhibition by KIRA6 or a pharmaceutical agent derived from KIRA6 as a pharmacophore.

Whereas LYN deficiency results in enhanced cytokine production upon Ag treatment (Parravicini et al., 2002) (Hernandez-Hansen et al., 2004) (Miranda et al., 2016), no increase has been observed in Ag-stimulated WT BMMCs under KIRA6 treatment. Though 1 µM KIRA6 appears to be required to fully inhibit FYN action, suboptimal suppression of FYN at lower concentrations of KIRA6 might already alleviate the potential cytokine-enhancing effect of LYN inhibition by KIRA6, resulting in a reduction of IL-6 and TNF production. Moreover, in addition to LYN and FYN, other SFKs (e.g. HCK and SRC) might also be slightly inhibited, and the composite effect could lead to attenuated cytokine production. In principle, SFKs can be classified into two subfamilies, where HCK can be regarded as LYN-related, and FYN as well as SRC belong to the SRC-related subfamily (Ingley, 2008).

Due to the structural similarities of SFKs, the generation of selective inhibitors for the particular kinases seems rather difficult. In addition, MCs express several different SFKs, namely LYN, FYN, SRC, and HCK (Suppl. Fig. 6B). Complicating the picture even more, most of them exert important, sometimes opposing functions (exemplified by Hong et al. (Hong et al., 2007), Parravicini et al. (Parravicini et al., 2002) and various publications on the role of LYN (Parravicini et al., 2002)(HernandezLJHansen et al., 2004) (Miranda et al., 2016). The frequently used inhibitors PP1 and PP2 are called SRC-family selective Tyr kinase inhibitors, which points out that these drugs are non-selective for the specific/individual SFK members. Moreover, the SFK least inhibited by these drugs, showing the highest IC_50_, appears to be LYN. Another well-known SFK inhibitor, SU6656, inhibits LYN with a fairly low IC_50_ (app. 130 nM). However, the respective IC_50_ for FYN unfortunately is very close (app. 170 nM), making this drug rather useless for a differentiation between LYN- and FYN-mediated functions. Our data reported in this study suggest a considerable difference concerning the concentrations of KIRA6 required for the inhibition of LYN and FYN. This suggests that the structure of KIRA6, respectively the relevant sub-structure/pharmacophore might be used as scaffold for the development of novel LYN- and FYN-selective inhibitors.

KIRA6 was developed as a kinase-inhibiting RNase-attenuating molecule (Ghosh et al., 2014). It is a type II inhibitor inhibiting IRE1α phosphorylation in an ATP-competitive manner stabilizing its inactive kinase conformation (Ghosh et al., 2014). Effectivity and selectivity were verified by performing ON-target competition tests, particularly with respect to IRE1α-mediated *XBP1* cleavage *in vitro* and *XBP1* mRNA splicing *in vivo*. It was further proven that KIRA6 does not inhibit kinase activity *in vitro* regarding a panel of Ser/Thr kinases (including ERK2, JNK2, and PKA) (Ghosh et al., 2014)). A further study applying kinase photoaffinity labeling combined with mass spectrometry analysis, however, suggested that KIRA6 might be not as selective as initially aimed for with the majority of off-targets not being protein kinases but mostly nucleotide-binding proteins (Korovesis et al., 2020)(Rufo et al., 2022). However, these analyses were performed in cellular lysates using a rather high concentration of labeled KIRA6 (10 µM) and thus it remains to be shown if such concentration can also be reached in living cells or if these proteins also serve as KIRA6 targets in living cells. Obviously, the highest concentration of KIRA6 needed in our study, depending on MC type studied, was 1 µM and the comparison of wild-type and LYN-deficient BMMCs clearly showed the necessity of presence of LYN for a KIRA6 effect in the nanomolar range. Intriguingly, though the MC-relevant kinases attenuated by KIRA6 in our present and former studies (Mahameed et al., 2019) are Tyr kinases, our modeling approach indicated that KIRA6 acts as type II inhibitor as well.

When deciding on the preferable quality of a pharmacological inhibitor in a situation-dependent manner, questions arise like i) how selective does an inhibitor has to be, and ii) how much promiscuity for substrate proteins can be allowed or even is desirable? In chronic lymphocytic leukemia, for instance, LYN has been demonstrated to be essential for the formation of a microenvironment supporting leukemic growth (Nguyen et al., 2016), and hence in such a situation, a high selectivity for LYN-inhibition would be desirable. On the other hand, in MCs upon IgE- or KIT-mediated activation, LYN represents an important activating signaling element with attenuating functions as well, hence potentially restricting application of highly selective inhibitors (Xiao et al., 2005)(Miranda et al., 2016)(Hernandez-Hansen et al., 2004). LYN-deficiency results in enhanced Ag-triggered pro-inflammatory cytokine production as well as in increased SCF-induced proliferation. Thus, for MC inhibition in the context of IgE-dependent allergic or KIT-driven neoplastic diseases, co-inhibition of LYN, FYN, and KIT would be an evident advantage. Hence, our molecular, cellular, as well as molecular modeling data will allow for the structure-guided design of novel inhibitors either selectively targeting single kinases or being highly specific for combinations of LYN, FYN, and/or KIT on the basis of KIRA6 as a pharmacophore.

## Acknowledgements

K. Maschke-Neuß is acknowledged for excellent technical assistance.

## Authorś Contributions

Conceptualization M.H., T.W.; Data curation V.W., T.W., G.R., J.G., M.A.M.; Formal analysis V.W., T.W., M.A.M., J.G., T.S.; Funding acquisition M.H., B.T., F.L.S.; Investigation VW, T.W., S.B., R.S., S.C., M.S., N.A., G.B., C.M. T.S..; Methodology G.R., J.G., M.A.M.; Project administration M.H., T.W.; Resources M.H., G.R., F.L.S., B.T., M.A.; Software G.R., M.A.M., J.G.; Supervision T.W., M.H., B.T., F.L.S., G.R.; Validation V.W, T.W.; Visualization VW, T.W., M.H.; Writing - original draft T.W., M.H., V.W.; Writing review & editing T.W., M.H., B.T., F.L.S., S.C., M.A., T.S..

