## Supplementary material for "KIRA6 is an effective and versatile mast cell inhibitor of IgE-mediated activation": Table 1

| Source | ID | Protein | State | ΔH/ kcal mol$^{-1}$ | ΔG/ kcal mol$^{-1}$ | Glide SP Docking Score* |
|---|---|---|---|---|---|---|
| Modeler | ---- | LYN | inactive/DFG-out | -60.6 ± 0.1 | -49.4 ± 3 | -14.4 |
| Modeler | ---- | FYN | inactive/DFG-out | -57 ± 0.1 | -33.5 ± 4.4 | -12.7 |
| XRAY | 6MOB | KIT | inactive/DFG-out | -67.2 ± 0.1 | -58.5 ± 3.2 | -15.0 |
| XRAY | 2DQ7 | FYN | active/DFG-in | -26.5 ± 0.2 | 4.9 ± -5.1 | -6.5 |
| XRAY | 5XY1 | LYN | active/DFG-in | -34 ± 0.1 | 1.3 ± 5.7 | -4.6 |
| XRAY | 1PKG | KIT | active/DFG-in | -27.1 ± 0.1 | -0.6 ± 3.5 | -5.5 |

Wunderle et al.
