## Supplementary figures and images for "KIRA6 is an effective and versatile mast cell inhibitor of IgE-mediated activation"

### Suppl. Fig. 1

#### Suppl. Fig. 1

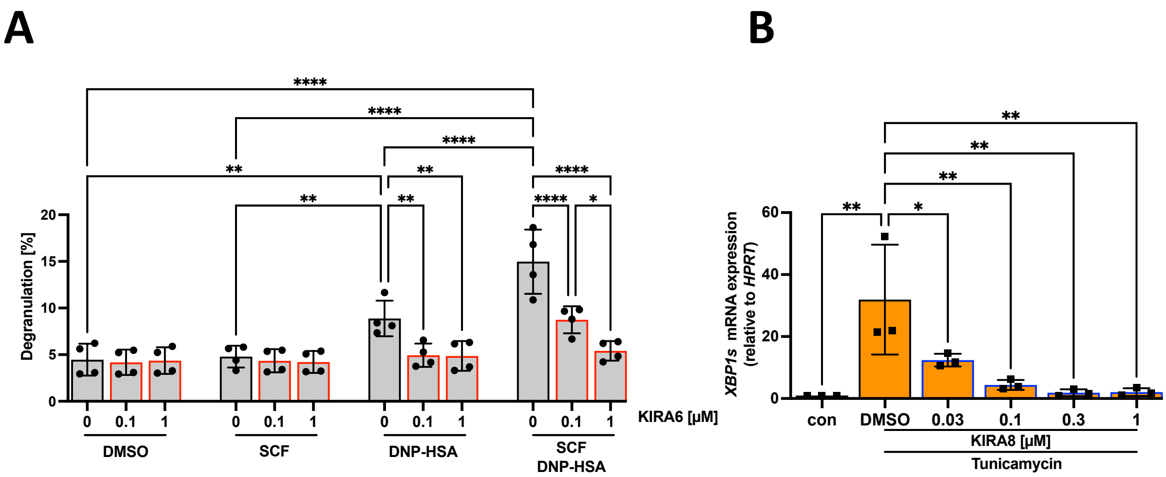


Wunderle et al.

### Suppl. Fig. 2

**Suppl. Fig. 2**

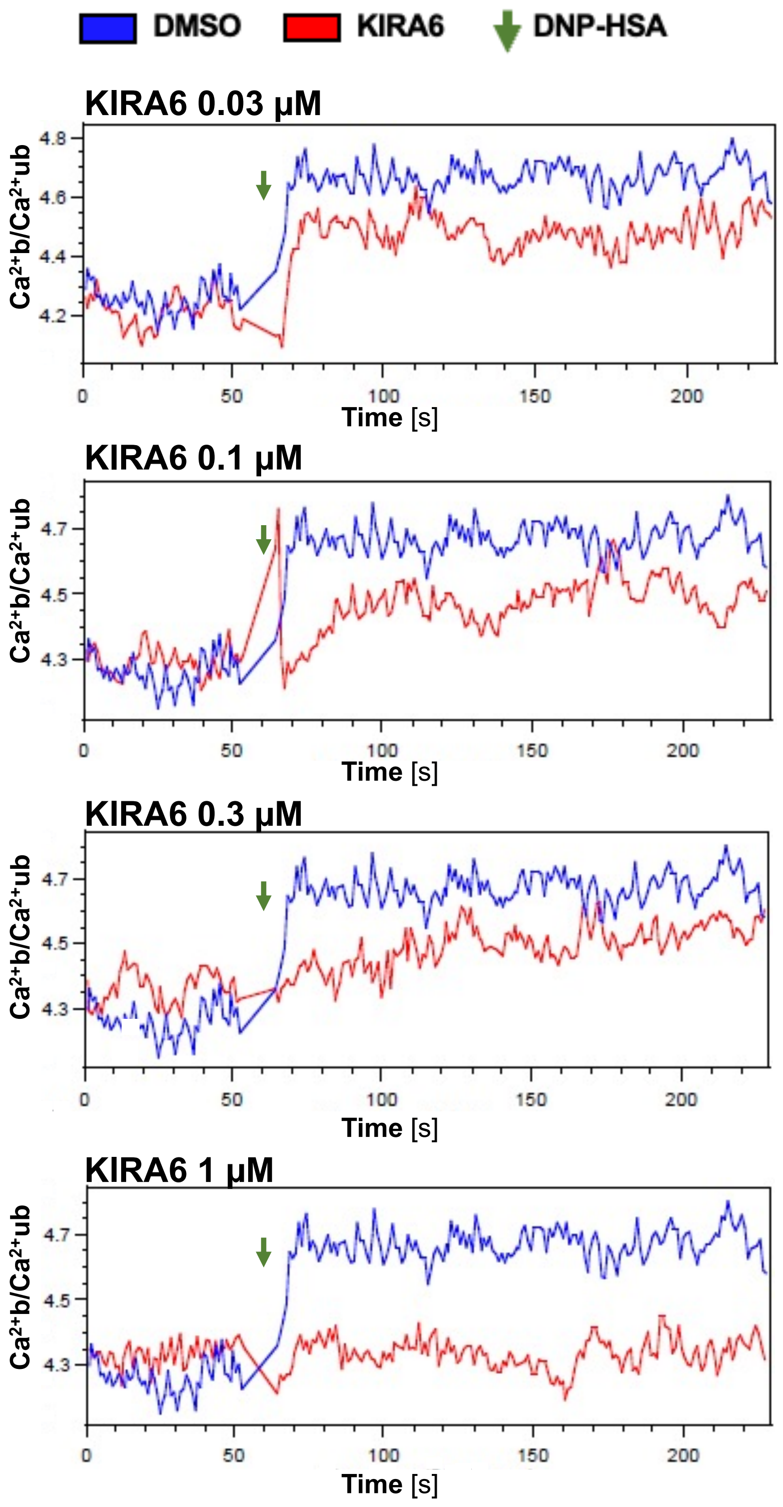


Wunderle et al.

### Suppl. Fig. 3

##### Suppl. Fig. 3

**A**

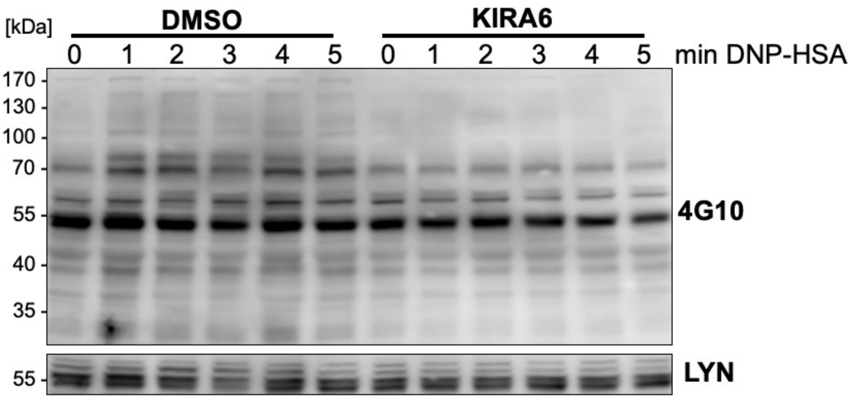


**B**

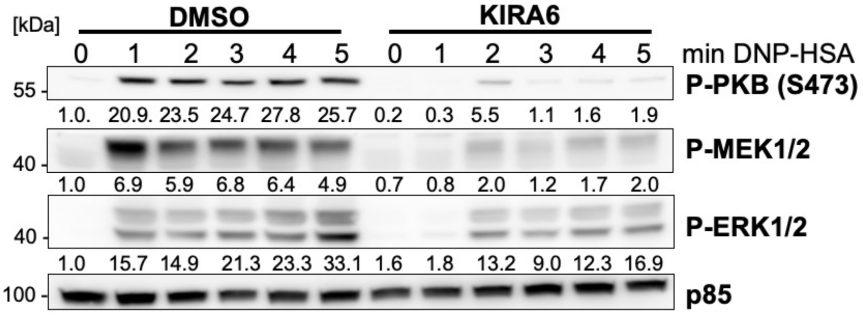


Wunderle et al.

### Suppl. Fig. 4

#### Suppl. Fig. 4

## A

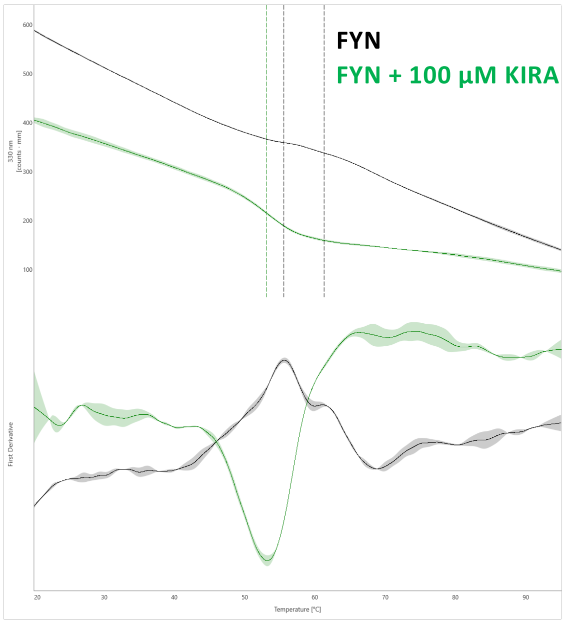


IP1 (FYN) = 55.6°C
IP1 (FYN+KIRA6) = 52.6°C
∆IP1 = -3.0°C
IP2 (FYN) = 61.4°C

## B

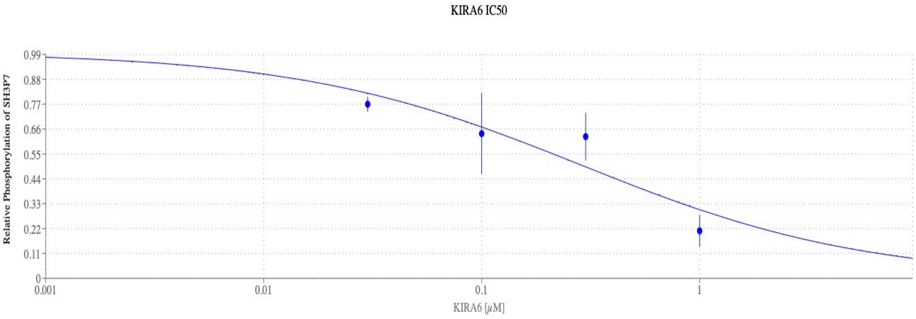


IC50 0.2876 µM

Wunderle et al.

### Suppl. Fig. 5

Suppl. Fig. 5

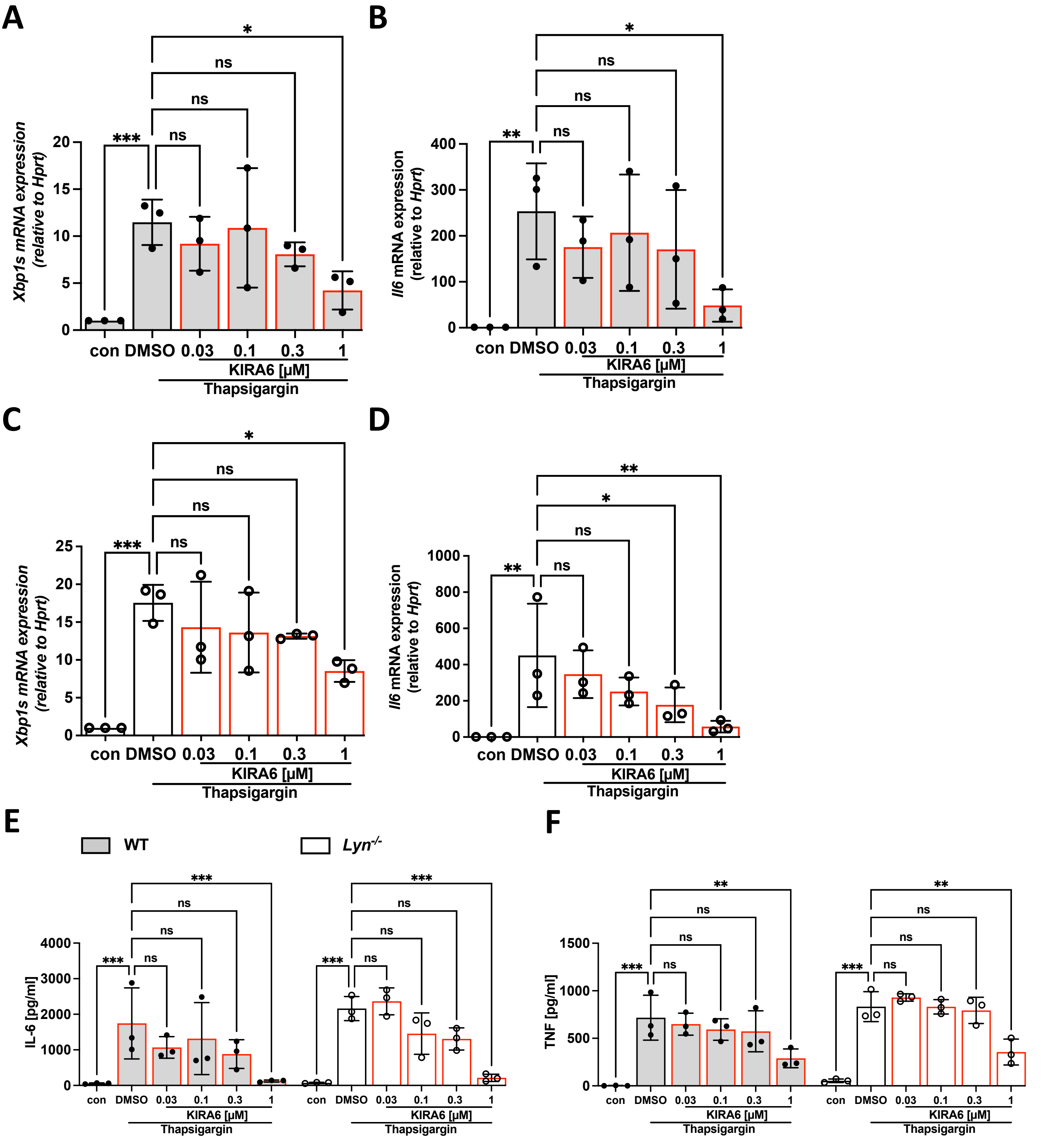


Wunderle et al.

### Suppl. Fig. 6

##### Suppl. Fig. 6

**A**

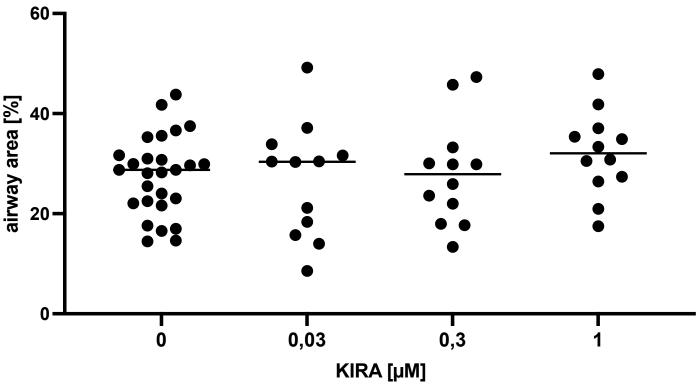


**B**

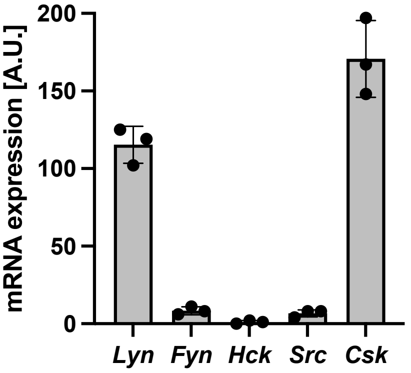


Wunderle et al.
